# Cellular composition of tissue-resident monocyte-lineage cells reveal functional heterogeneity during inflammatory arthritis

**DOI:** 10.1101/2025.06.13.659432

**Authors:** Yidan Wang, Samuel D. Dowling, Jessica Maciuch, Vanessa Rodriguez, Meghan Mayer, Kainat Mian, Tyler Therron, Hadijat-Kubura M. Makinde, Carla M. Cuda, Deborah R. Winter, Harris Perlman

## Abstract

Tissue-resident monocyte-lineage cell (TRMC) are an extravascular population distinct from circulating monocytes and synovial macrophages and are critical for the development of inflammatory arthritis. However, the precise identities and origins of TRMC subpopulations remain unclear. Here, we characterize the ontogeny of TRMC, which are comprised of bone-marrow (BM)-derived and an embryonic, long-lived population. Furthermore, we identified three TRMC subpopulations, distinguished by expression of TIM4, CX3CR1, and MHCII. Clodronate-laden liposome reduces the number of TRMC but does not impact the proportions or transcriptional profile of TRMC subpopulations at 7 days post administration. TIM4^+^CX3CR1^+^ and TIM4^+^ TRMC represent long-lived population, whereas MHCII+ TRMC are BM-derived and dependent on Ccr2 during steady state. BM-derived TRMC expand and replenish the TIM4^+^CX3CR1^+^ and TIM4^+^ TRMC compartments throughout the peak and plateau of inflammatory arthritis. These findings underscore the importance heterogeneity within TRMC and highlight their distinct responses to synovial disruption and potential roles in rheumatoid arthritis (RA).

## INTRODUCTION

Rheumatoid arthritis (RA) is a systemic autoimmune disease, characterized by leukocyte trafficking, cartilage destruction, pannus formation and bone erosion ^1–4^. One of the major cell types responsible for persistent inflammation in the synovium are monocytes. To model the effector phase of RA, researchers commonly use the K/BxN serum transfer-induced arthritis (STIA) mouse model ^5–8^. Pharmacologic depletion of all monocytes with liposomal clodronate completely prevents the development of arthritis ^6,7^. Furthermore, adoptive transfer of flow cytometry sorted non-classical monocytes could rescue arthritis in clodronate depleted animals ^7^. These data suggest that circulating non-classical monocytes (NCM) are essential for STIA. In contrast, mice genetically deficient in classical monocytes (Ccr2^−/−^) or NCM (Nr4a1^−/−^ and Nr4a1 super enhancer domain 2 (SE2)-deficient) as well as Ccr2^−/−^Nr4a1^SE2-def^ double knock-out mice still develop RA-like disease ^6^. Collectively, these data suggest a previously unidentified population of monocytes is necessary and sufficient for the development of arthritis.

Monocyte-lineage cells are a heterogeneous population in peripheral blood (PB) and synovial tissue. There are three major monocyte subsets in PB of mice: classical monocytes (CM), characterized as CCR2^+^Ly6C^Hi^CD62L^+^CX3CR1^Low^CD43^−^; intermediate monocytes (IM), defined as CCR2^−^Ly6C^Int^CD62L^−^CX3CR1^int^CD43^±^; and non-classical monocytes (NCM), identified as CCR2^−^Ly6C^Low^CD62L^−^CX3CR1^Hi^CD43^+^ ^9–11^. Studies on monocyte-lineage cells focused on circulating cells, while only a few studies have investigated monocyte-lineage cells in tissues ^6,12–14^. Silva H, *et al.* reported the existence of Ly6C^+^ monocytes in adipose tissue that is transcriptionally distinct from blood monocytes and macrophages ^12^. Similarly, a Ly6C^−^ monocyte population was identified in lung though this population was highly labeled by CD45 intravascular labeling (I.V.) ^13^. However, the tissue-resident monocytes in both studies express CD64, a known marker for macrophages, raising questions about whether it represents a separate macrophage subset ^12^. Another study showed that muramyl dipeptide (MDP) induced Nod2-dependent, Ccr2-dependent, Nr4a1-independent Ly6c^low^ NCM population (I-NCM) infiltrates tumors, resembles macrophages and dendritic cells transcriptionally, and is essential for reducing lung tumor metastasis ^14^. Although I-NCM represents a true monocyte population, they do not exist during homeostasis ^14^.

In synovial tissue, monocyte-lineage cells include intra- and extravascular CD64^−^Ly6C^−^ compartments ^6^. The intravascular CD64⁻Ly6C⁻ cells resemble circulating non-classical monocytes (NCM), while the extravascular CD64⁻Ly6C⁻ cells represent a newly identified synovial monocyte-lineage population, termed tissue-resident monocyte-lineage cells (TRMC) ^6^. TRMC are the only reported true monocyte-lineage cells that reside in tissue during steady state. TRMC lack the macrophage surface marker CD64, and are transcriptionally different from circulating monocytes, synovial dendritic cells (DC), and macrophages. Unlike macrophages, TRMC are also sensitive to depletion by clodronate-laden liposome. In response to synovial inflammation, TRMC respond by expanding and enriching the pro-inflammatory associated pathways. TRMC are long-lived and embryonically derived using CX3CR1^CreER^.zsGFP fate-mapping mice, supporting the different ontogenies of TRMC and the possible existence of heterogenic subpopulations. However, the precise identities of TRMC subpopulations and whether they respond differentially to synovial disruption or inflammation remain unknown.

Here, we investigated TRMC heterogeneity and function across with a focus on their monocyte-derived vs. tissue-resident origin. We identified three TRMC subpopulations with distinct functions during steady state: TIM4^+^CX3CR1^+^, TIM4^+^, and MHCII^+^ TRMC, which originate from two distinct sources: long-lived and bone-marrow (BM) derived TRMC. Notably, only long-lived TRMC are sensitive to Clo-lip depletion, whereas the survival of BM derived TRMC is Ccr2-dependent. During K/BxN serum-transfer-induced arthritis (STIA), the number of BM-derived TRMC exceeded that of long-lived TR-MC, in contrast to the steady state where long-lived TRMC predominate. These results highlight the heterogeneity of TR-MC and their central roles in arthritis development.

## METHODS

### Mice

All mice used in the study were females and aged between 8-12 weeks or 8-20 weeks post-bone marrow restoration of bone marrow chimeras (BMC) mice. All strains of mice were on C56BL/6 (B6) background with specific details listed in experimental mouse strain table. All mice were bred and housed in the barrier animal facility of Northwestern University. All experimental procedures on mice reported in the study were approved by Institutional Animal Care and Use Committee (IACUC) office.

### Bone marrow chimeras

Starting one week prior to bone marrow chimeras (BMC) generation and continuing for 5 weeks afterwards, host mice including CD45.1 B6, CD45.2 B6, and Ccr2^−/−^ mice were placed on water bottles containing Sulfamethoxazole and Trimethoprim (Novitium Pharma, 70954-258-10). One hundred milligram (mg)/1 kilo gram (kg) mouse weight Ketamine (Dechra, ANADA 200-073) and 20mg/kg xylazine (AnaSed Injection, NDC 59399-110-20) were diluted in sterile WFI water to a total volume of 100 microliters (µL) and used for anesthesia of the host mice right before irradiation. One thousand centigray (cGy) γ irradiation was performed on each host mouse. Six hours post the irradiation procedure, 40mg/kg busulfan (Caymen Chemical, 55-98-1) was administered intraperitoneally (IP). A single-cell suspension from the bone marrow of donor mice with different CD45 allotype, containing a minimum of 10 million cells, was administered retro-orbitally to the host mice 24 hours post the irradiation process. To determine the success of chimerism, peripheral blood (PB) was collected from the submandibular vein 6-8 weeks after the bone marrow cells transfer.

### Parabiosis

The generation of parabiotic mice was performed as described in Kamaran P, *et al.* ^15^ between pairs of mice with different CD45 allotypes and conducted at Microsurgery Core, Northwestern University. Starting one week prior to the parabiosis surgery and 5 weeks afterwards, mice were placed on water bottles containing Sulfamethoxazole and Trimethoprim (Novitium Pharma, 70954-258-10). Parabiotic mice received daily subcutaneous sterile saline (Teknova, S5820) injections for two weeks following surgery as post-operative care.

### Single-cell suspension preparation

To prepare single-cell suspension from PB, PB was first withdrawn via either heart stick or submandibular vein and processed as previously described ^6,7^. To prevent non-specific binding, 0.5mg CD16/32 Fc Block (BD Biosciences, 553142) was used to incubate the cells at 4℃ for 20 minutes. Antibody cocktail (R718 CD45.1 (BD Biosciences, 567297), BV421 CD45.2 (BioLegend, 109831), BV711 Ly6G (BD Biosciences, 563979), PE-CF594 SiglecF (BD Biosciences, 562757), BB700 CD11b (BD Biosciences, 566416), APC CD4 (BD Biosciences, 553051), APC CD8a (BD Biosciences, 553035), APC CD19 (BD Biosciences, 550992), APC NK1.1 (BD Biosciences, 550627), PE CD115 (Invitrogen, 12-1152-82), BV650 CX3CR1 (Biolegend, 149033), PE Cy7 CD62L (BioLegend, 104418), APC Cy7 (BD Biosciences, 560596)) was then added for an incubation of 30 minutes at 4℃. Erythrocytes lysis and tissue fixation were performed using FACS Lyse (BD Biosciencess, 349202) at room temperature for 10 minutes.

As previously described ^6^, intravascular cells were labeled using 1ug anti-CD45-BUV661 antibody (BD Biosciences, 612975), administered retro-orbitally to the mouse and incubated for 4 minutes with mouse being active during the time window. To prepare a synovial (Syn.) single-cell suspension from mouse tissue, the ankles were dissected by cutting 3 millimeters (mm) above the hind paws. The skin and toes of the ankles were removed, leaving the knuckles intact. Bone marrow of each ankle was flushed using 2.5 mL HBSS (Gibco, 14025092). Digestion buffer, consisting of 2.4mg/mL dispase II (Roche, 4942078001), 2mg/mL collagenase D (Roche, 11088866001), 0.2mg/mL DNAse I (Roche, 10104159001) in HBSS with pH adjust to 7.2-7.4, was used to influx the synovium via the posterior aspect of the tibiotalar joint. Ankles were incubated for 1 hour at 200 rpm and 37℃. The incubation was suspended with autoMACS^TM^ Running Buffer (Miltenyi Biotec, 130-091-221). The ankles were then gently smashed through 40mm nylon cell strainers (FALCON, 352340) for a total of three times. Erythrocytes were lysed using 200mL 1 X BD Pharm Lyse lysing buffer (BD Biosciences, 555899) for 1 minute. Dead cells were stained using eFluor^TM^ 506 Fixable Viability Dye (Invitrogen, 65-0886-14) for 15 minutes. Per 5 millions of cells, 0.5mg CD16/32 Fc Block (BD Biosciences, 553142) was used to incubate the cells at 4 ℃ for 20 minutes, followed by antibody staining (BUV395 CD45 (BD Biosciences, 565967), R718 CD45.1 (BD Biosciences, 567297), BV421 CD45.2 (BioLegend, 109831), BB700 CD11b (BD Biosciences, 566416), BV711 Ly6G (BD Biosciences, 563979), PE-CF594 SiglecF (BD Biosciences, 562757), BV786 CD64 (BD Biosciences, 741024), PE Cy7 MHCII (BioLegend, 107630), APC Cy7 (BD Biosciences, 560596), AF647 CD177 (BD Biosciences, 566599), BUV737 Treml4 (BD Biosciences, 755275), BV650 CX3CR1 (Biolegend, 149033)) at 4℃ for 30 minutes. Fixation of cells was performed at room temperature in dark for 15 minutes using 2% paraformaldehyde solution (Electron Microscopy Sciences, 15713-S) diluted in in PBS.

### Flow cytometry analysis

All data were obtained at Robert H. Lurie Comprehensive Cancer Center (RHLCCC) Flow Cytometry Core Facility, Northwestern University. Analytical flow cytometry data were acquired on Symphony A5.2 Spectral Analyzer (BD Biosciences). Florescence activated cell sorting (FACS) was performed on FACS^TM^ III Cell Sorter (BD Biosciences). When necessary, fluorescence minus one (FMO) was applied to set gating thresholds. Absolute cell numbers were calculated by adding 123count eBeads^TM^ Counting Beads (Invitrogen, 01-1234-42). Compensation calculation and unmixing as well as analysis were all conducted using FlowJo (v10.7.1)

### K/BxN serum transfer induced arthritis (STIA)

The serum from K/BxN transgenic mice was titrated by assessing the clinical scores of the ankles every 2–4 days using the following scoring system: 0 = no inflammation; 1 = swollen toes with the ankle maintaining a funnel shape; 2 = swollen ankle with the funnel shape disappearing; 3 = swollen ankle with an inverted funnel shape. A titrated serum was selected when the sum of scores from all four ankles of a mouse reached approximately 10 at the peak of the disease, sustained for around 5 days during the plateau stage, and gradually resolved 28–32 days post-administration. The titrated serum from K/BxN transgenic mice was administered retro-orbitally to recipient mice. STIA development was induced independently with titrated K/BxN serum for flow cytometry and cellular indexing of transcriptomes and epitopes by sequencing (CITE-seq).

### Clodronate liposome (Clo-lip) administration

For Clo-lip administration, 200 µL of Clo-lip was injected retro-orbitally into BMC mice. To confirm successful depletion, peripheral blood (PB) was collected via the submandibular vein, and monocyte levels in PB were assessed by flow at days 0, 1, 3, 7 following Clo-lip administration. Synovial tissue for flow cytometry and CITE-seq was collected from independent mice cohorts treated with Clo-lip in parallel.

### Cellular Indexing of Transcriptomes and Epitopes by sequencing (CITE-seq)

For all CITE-seq studies, live CD45^+^CD11b^+^Ly6G^−^SiglecF^−^MHCII^low/mid^CD64^−^CD177^−^ monocyte-lineage Syn. cells from CD45.1→CD45.2 BMC mice were sorted from a single cell suspension as described above. Generation of GEMs, barcoding, cDNA amplification, 3′ gene libraries, and cell surface protein libraries was conducted using the Chromium Next GEM Single Cell 3′ (v3.1, dual index) protocol with Feature Barcode technology at the Metabolomics Core, Northwestern University. Sequencing was performed on a NextSeq 2000 (Illumina) at the same facility. The mouse reference genome mm10 (mm10-2020-A) was used for read alignment and quantification through the count function of the Cell Ranger pipeline (v6.1.2). Seurat (v4.4.0) package was utilized for quality control, preprocessing, clustering, identification of *de novo* markers, label transfer, and visualization. Quality control (QC) for each dataset was assessed based on UMI counts, mitochondrial gene percentages, and ribosomal gene expression and thresholds were set as defined below. Doublets were identified and removed using scDblFinder (v1.12.0) with an expected doublet formation rate of 5%. Detailed QC filtering information for each sample is provided below.

| Genotype | # Cells | Median | Median | Min | Max | Max | Max | # Cells |
| --- | --- | --- | --- | --- | --- | --- | --- | --- |
|  | Detected | Genes/ | UMIs/ | UMI/ | UMI/ | % MT | % Rb | After |
|  |  | Cell | Cell | Cell | Cell | reads/ | reads/ | Filtering |
|  |  |  |  |  |  | cell | cell |  |
| Steady state | 1912 | 2695 | 8562 | 3500 | 40000 | 10 | 25 | 1273 |
| Day 7 Clo-lip | 1277 | 2548 | 7967 | 3500 | 40000 | 10 | 25 | 861 |
| Baseline STIA | 1128 | 2283 | 7434 | 3500 | 35000 | 10 | 25 | 678 |
| Peak STIA | 6490 | 2226 | 7939 | 3500 | 35000 | 10 | 25 | 4382 |
| Plateau STIA | 3104 | 2119 | 6450 | 3500 | 35000 | 10 | 25 | 1834 |
| Resolution STIA | 4779 | 1803 | 4771 | 3500 | 35000 | 10 | 25 | 2856 |

\* Mitochondrial - MT; ribosomal - Rb

For the steady state BMC dataset, SCTransform was performed using 750 variable genes for dimensionality reduction and clustering. The top 10 PCs were selected for generating the UMAP and constructing a shared nearest neighbor graph using FindNeighbors (with 10 nearest neighbors), followed by clustering with FindClusters (resolution = 0.2). Antibody-derived tag (ADT) counts were normalized within each cell using the centered log-ratio (CLR) method to assess surface markers expression. Differentially expressed genes (DEG) identification and Pearson correlation analysis were performed under the RNA assay after initial log-normalization and subsequently scaling across 2,000 variable features with the ScaleData function. DEG of each cell types were performed using FindAllMarkers with logfc.threshold = 0.25 and min.pct = 0.25 using the Wilcoxon test with an adjusted p-value < 0.05, corrected by the Benjamini-Hochberg (BH) method. Cluster annotation was determined using canonical gene expression, ADT expression, and de novo marker genes identified by the FindAllMarkers function. Pearson correlation analysis was performed on the aggregated gene expression data for each annotated cell type under the RNA assay.

The Day 7 post Clo-lip BMC dataset, which was batched with the steady state sample, was similarly normalized using SCTransform on 750 variable genes and annotated by label transfer using FindTransferAnchors (18 PCs; 20 k.anchors) and TransferData (18 PCs) with the steady state BMC dataset as reference. The number of PCs was determined by selecting the minimum number at which the canonical genes of TRMC and monocytes showed prominent loadings. MapQuery was used for the projection of Day 7 post Clo-lip BMC dataset onto steady state BMC dataset UMAP structure.

Baseline, peak, plateau, and resolution STIA BMC datasets were preprocessed as mentioned above with 1000 genes selected as variable features for SCTransform. Label transfer was conducted using FindTransferAnchors (13 PC; 20 k.anchors) and TransferData (13 PC) for annotation with the Steady-state BMC data set as reference. MapQuery was used for the projection of 4 STIA BMC datasets onto steady state BMC dataset UMAP structure. Osteoclasts were identified by module score calculated using five manually selected canonical osteoclast genes (Dcstamp, Ctsk, Acp5, Mmp9, Oscar) with a threshold over 1.35 of osteoclast module score across the four STIA BMC datasets. For trajectory inference, classical monocytes (CM), non-classical monocytes (NCM), and the three TRMC subpopulations from all 4 STIA BMC datasets were merged and analyzed using Monocle 3 (v1.3.7). Normalization and dimension reduction were carried out using preprocess_cds with PC = 30 and default parameters. UMAP projection was performed using reduce_dimension function, with batch effect corrected using align_cds. A principal graph representing the differentiation trajectory was constructed using learn_graph with close_loop = FALSE. Genes exhibiting significant variation along the trajectory were identified using graph_test() and clustered into co-expressed gene modules using find_gene_modules().

The AddModuleScore function was used to calculate the module scores of TR-MC subsets in Day 7 post Clo-lip BMC dataset, baseline, peak, plateau, and resolution STIA BMC datasets based on the top 10 DEG ranked by the average log2 fold-change removing any overlapping DEGs across 3 TRMC subsets from steady state BMC dataset. Gene ontology (GO) enrichment analysis was conducted on for each cell type based on marker defined using FindAllMarkers with logfc.threshold = 0.25, only.pos = TRUE, and min.pct = 0.25, compared to a background genes defined using FindAllMarkers with logfc.threshold = 0, only.pos = FALSE, and min.pct = 0.25. This analysis was conducted using clusterProfiler package (v4.6.2) with a BH adjusted p-value < 0.05. Pseudobulk RNA-seq analysis of the aggregated RNA expression of the three TRMC subsets across the baseline, peak, plateau, and resolution STIA BMC datasets was conducted using AggregateExpression() function from the merged dataset. Gene expression values were normalized using varianceStabilizingTransformation() and subsequently scaling using scale(). K-means clustering was applied on the pseudobulk gene expression values with k = 5 using kmeans() function. Gene set variation analysis (GSVA) was calculated on gene ontology biological process (GO_BP) at single-cell level for each of the three TRMC subpopulations merged across the 4 STIA BMC datasets. Enriched pathways were filtered by selecting the pathways with the difference between the highest and lowest average enrichment score across cell types from each STIA dataset > 0.8 and containing at least 3 genes expressed in the merged dataset.

### Statistical analysis

For all experimental results, the numbers displayed in the figures are mean of all the datapoints. Unpaired t-test was performed for all comparisons of cell numbers, except for the comparison of cell origins distribution in parabiosis mice, which was calculated using a chi-square test. Statistical significance was defined as p-value less than 0.5, and all analyses were performed in GraphPad Prism (v10.3.0). Numbers of mice used for each study is specified on the plots or in the figure legends.

## RESULTS

### TRMC are a heterogenous cell population

To identify whether TRMC are a heterogeneous population, we generated CD45.1→CD45.2 C57BL/6 bone marrow chimeras (BMC) mice with their ankles shielded to preserve tissue-resident cells for the investigation of the heterogeneity of TRMC (Fig 1A). TRMC were defined as CD45^+^CD11b^+^Ly6G^−^SiglecF^−^MHCII^mid/low^CD64^−^Ly6C^−^CD177^−^I.V.CD45^−^, consistent with the previously described definition with the additional exclusion of Ly6C^low^CD177^+^ neutrophils ^6,16^ (Supplemental Fig 1A). Synovial (Syn.) classical monocytes (CM) and non-classical monocytes (NCM) were defined as CD45^+^CD11b^+^Ly6G^−^SiglecF^−^MHCII^mid/low^CD64^−^Ly6C^+^ and CD45^+^CD11b^+^Ly6G^−^SiglecF^−^MHCII^mid/low^CD64^−^Ly6C^−^CD177^−^I.V.CD45^+^, respectively (Supplemental Fig 1A). Almost 100% of PB CM and NCM were donor-derived cells, suggesting generation of successful and complete chimera for circulating monocytes (Supplemental Fig 1B). In synovial tissue, an average of 84.8% of Syn. CM and 97.9% of Syn. NCM were donor-derived cells, whilst Syn. TRMC were 61% host-derived and 39% donor-derived (Fig 1B-C). UMAP projection of Syn. CM, Syn. NCM, host-derived TRMC, and donor-derived TRMC showed a distinct distribution of the four cell types, suggesting their unique surface protein expression profiles based on a combination of surface markers excluding CD45.1 and CD45.2 (Live/Dead, CD45, CD11b, Ly6G, SiglecF, CD64, MHCII, CD177, Ly6C, I.V. CD45) (Supplemental Fig 1C). Next, we compared the expression patterns of 3 surface markers (TIM4, CX3CR1, MHCII) that have been reported as markers for tissue-resident or monocyte-derived macrophages ^7,8,17,18^. We observed a noticeably higher expression of TIM4 and a slightly higher expression of CX3CR1 in host-derived TRMC compared to donor-derived TRMC, whereas donor-derived TRMC exhibited higher MHCII expression than host-derived TRMC (Fig 1D). Although BMC mice are widely used to study the ontogeny and heterogeneity of cells, parabiosis is a better model for closely mimicking the steady state, with a nearly 1:1 chimera, since it avoids the trauma introduced by irradiation. (Fig 1E; Supplemental Fig 1D). Therefore, we also investigated the compartment of partner derived cells in CD45.1:CD45.2 parabiotic mice. Treml4 was sufficient to be used as a proxy for I.V. CD45 to identify TRMC (Supplemental Fig 1E). Less than 10% of PB and Syn. CM and ∼25% of PB and Syn. NCM were derived from the partner mouse (Fig 1G). A small percentage (∼4%) of TRMC were partner-derived cells (Fig 1F-G). Together, these results suggest TRMC is a heterogeneous population with distinct ontogenies.

**Figure 1.**
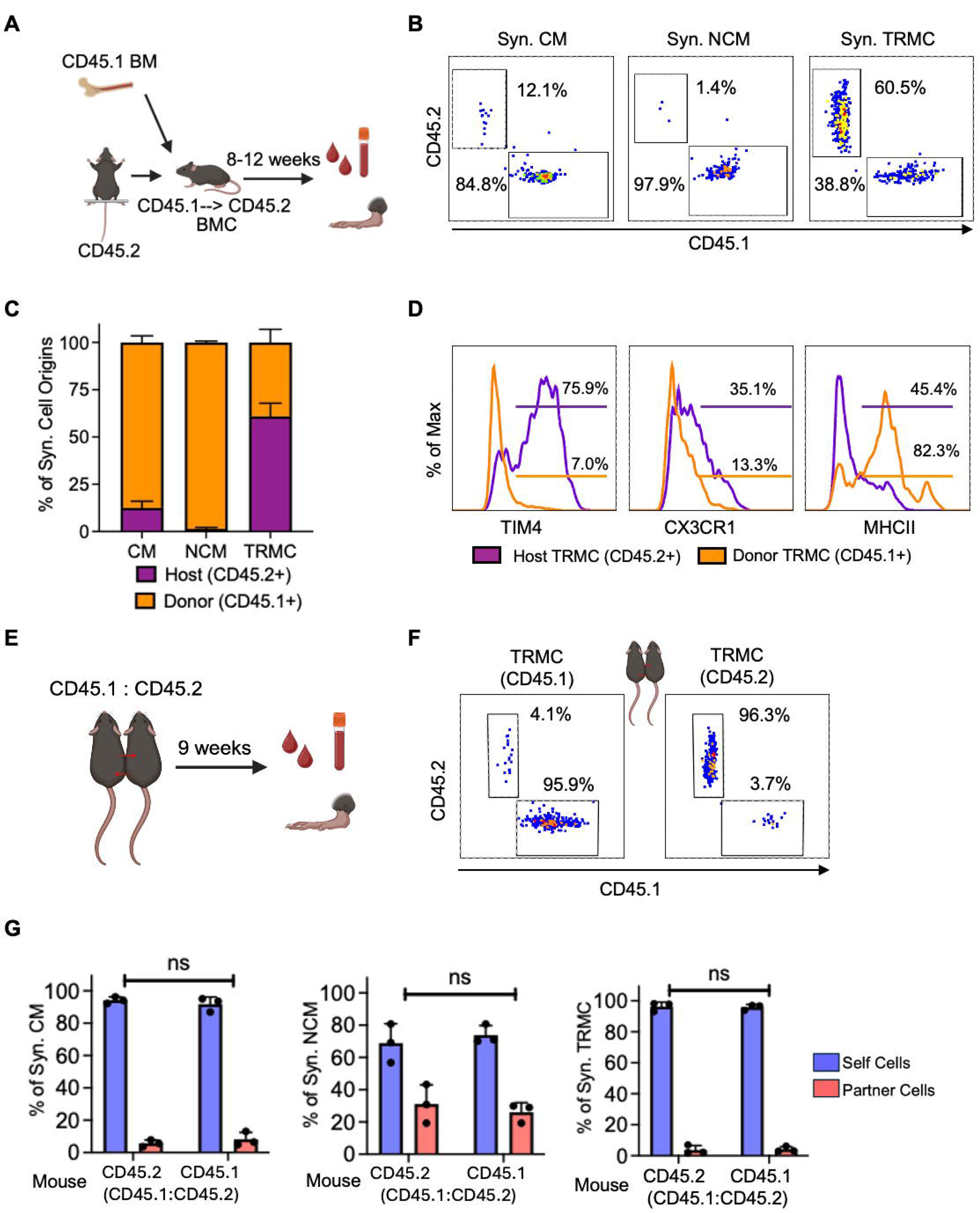
TRMC is a heterogeneous cell population. (A) Schematic mouse model of CD45.1 C57BL/6 (B6) to CD45.2 B6 bone marrow chimera mouse (CD45.1→CD45.2 BMC). (B) Expression of CD45.1 and CD45.2 in synovial (Syn.) classical monocytes (CM), Syn. Non-classical monocytes (NCM), and Syn. tissue-resident monocyte-lineage cells (TRMC) in steady state CD45.1→CD45.2 BMC (numbers are representatives of the averages of n= 6). (C) Percent of host-derived and donor-derived Syn. CM, Syn. NCM, and Syn. TRMC in steady state CD45.1→CD45.2 BMC (n= 6). (D) Expression of TIM4, CX3CR1, and MHCII in host-derived and donor-derived TRMC. (E) Schematic mouse model of CD45.1:CD45.2 parabiotic mouse model. (F) Expression of CD45.1 and CD45.2 in Syn. TRMC of CD45.1 B6 and CD45.2 B6 from CD45.1:CD45.2 parabiotic mice (numbers indicate the mean of the experiment n =3; Treml4 was used as a proxy for I.V. CD45 to identify TRMC). (G) Percents of self and partner cells of Syn. CM, NCM, and TRMC in CD45.1:CD45.2 B6 parabiotic mice (n = 3). Ns indicates p-value > 0.05 (chi-square test). All graphs are displayed as n ± SD.

### TRMC exists as subpopulations

To validate and further investigate the heterogeneity of TRMC, we performed CITE-seq, which provides both RNA transcriptional and antibody-derived tags (ADT) surface marker information, on FACsorted CD45^+^CD11b^+^Ly6G^−^SiglecF^−^MHCII^mid/low^CD177^−^ synovial cells in steady state from CD45.1→CD45.2 BMC mice. Based on canonical and de novo cell type markers, we identified distinct cell types including TRMC, classical monocytes, non-classical monocytes, cycling cells, and fibroblasts, which escaped from the sorting process (Fig 2A-D; Supplemental Fig 2A-D; Supplemental Table 1A). The intensity of ADT markers Tim4, CX3CR1, I-A-I-E (MHCII) distinguished three TRMC subpopulations as TIM4^+^CX3CR1^+^ TRMC, TIM4^+^ TRMC, and MHCII^+^ TRMC (Fig 2D; Supplemental Table 1B). The majority of TRMC during steady state were TIM4^+^CX3CR1^+^ TRMC (45.6%), followed by TIM4^+^ TRMC (40.3%), while MHCII^+^ TRMC constituted a minor population (14.1%) (Fig 2E). We annotated all cells as CD45.1^+^, CD45.2^+^, and double negative or ambiguous cells based on the ADT expression of CD45.1 and CD45.2 (Fig 2F; Supplemental Fig 2E-F). TIM4^+^CX3CR1^+^ TRMC and the majority of TIM4^+^ TRMC were CD45.2^+^, host-derived (Fig 2G). In contrast, nearly all MHCII^+^ TRMC were CD45.1^+^, donor-derived (Fig 2G). Comparison of the three TRMC subpopulations revealed distinct functional enrichments (Fig 2H; Supplemental Table 1C; Supplemental Table 2A-C). TIM4⁺CX3CR1⁺ TRMCs were enriched in genes associated with bone and muscle tissue development, including Sparc, which encodes protein involved in collagen calcification ^19^, and S100b, a calcium-binding protein essential for muscle contraction ^20^. Both TIM4⁺CX3CR1⁺ TRMC and TIM4⁺ TRMC were enriched in positive regulation of macrophage migration pathways, with differential expressed genes (DEG) such as Csf1r in TIM4⁺CX3CR1⁺ TRMC, which is involved in macrophage motility ^21^, and Ccl2 in TIM4⁺ TRMC, which is a component of the chemokine system ^22^. TIM4⁺ TRMCs show additional enrichment in chemokine-mediated signaling pathways, with genes such as Ccl24 ^23^, and the cellular response to hypoxia, featuring Pgk1, coding for enzyme required for glycolysis ^24,25^. MHCII⁺ TRMCs similarly exhibited enrichment in the cellular response to hypoxia as TIM4⁺ TRMCs and were further enriched in peptide antigen assembly with the MHC protein complex (H2-Aa and H2-Eb1) and regulation of cellular extravasation pathways, with key genes such as Ccr2 ^26,27^. These data demonstrate that TRMC exist as three functionally distinct subpopulations, each characterized by unique transcriptional and surface markers.

**Figure. 2.**
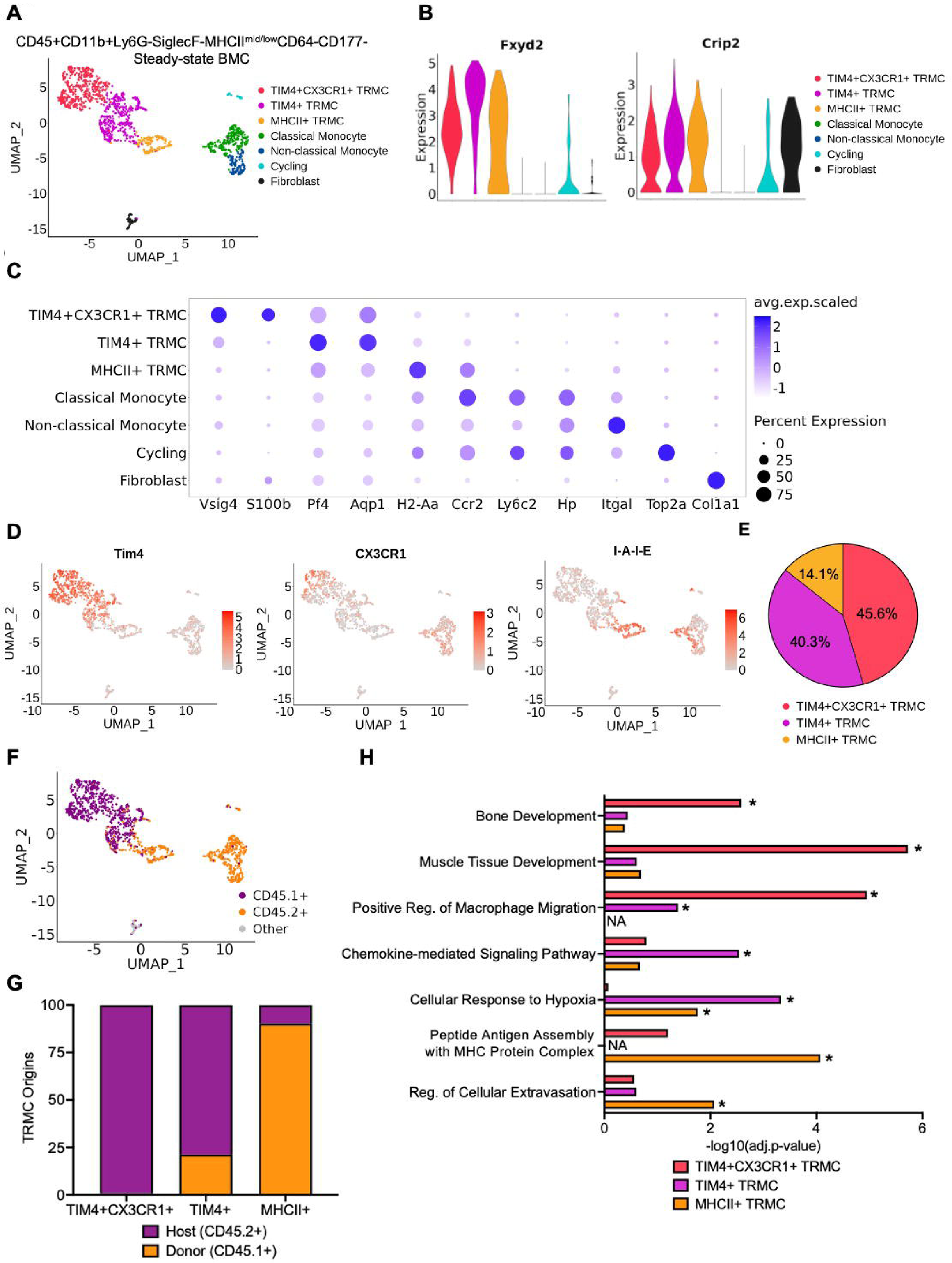
CITE-seq analysis identified distinct TRMC subpopulations. (A) Annotation of the seven synovial subpopulations in steady-state CD45.1 to CD45.2 BMC CD45^+^CD11b^+^Ly6G^−^ SiglecF^−^MHCII^mid/low^CD64^−^CD177^−^ cells displayed as Uniform manifold approximation and projection (UMAP). (B) Violin plots of TRMC canonical markers Fxyd2 and Crip2. (C) Bubble plots of canonical markers for TRMC subsets, CM, NCM, Cycling cells, and Fibroblast. (D) Feature plots of ADT expression of Tim4 (TIM4), CX3CR1, I-A-I-E (MHCII). (E) Proportion of the three subpopulations within TRMC. (F) Annotation of CD45.1+ and CD45.2+ cells. (G) Origins of donor-derived (CD45.1+) and host-derived (CD45.2+) cells of the three TRMC subpopulations. (H) Enriched gene ontology (GO) pathways in three TRMC subpopulations. Regulation (Reg.). Adjusted p values were calculated using Wilcox single-rank test and corrected using the Benjamin-Hochberg (BH) procedure. * Adjust p < 0.05.

### TRMC remain transcriptionally comparable after clodronate liposome (Clo-lip) treatment

We previously reported that TRMC are sensitive to Clo-lip, however, it remains unclear which TRMC subpopulations are affected and whether their transcriptional profiles are altered by Clo-lip. We administered Clo-lip. and measured the number of host-derived and donor derived cells in CD45.1→CD45.2 BMC mice over a seven-day period (Supplemental Fig 3A). As expected, donor-derived CM and NCM in both PB and synovial tissue were almost completely depleted one day post Clo-lip treatment and restored to the level comparable to steady state by day seven post treatment. A similar pattern was also observed in host-derived CM and NCM in both PB and synovial tissue (Supplemental Fig 3B-E). Host-derived synovial TRMC were significantly reduced on day 3 and day 7 post Clo-lip treatment, while changes in the donor-derived TRMC were not significant (Fig 3A). We observed a shift in the ratio of donor-derived to host-derived TRMC one day post Clo-lip, with donor-derived TRMC becoming the predominant population (Fig 3B). The finding is surprising, given the complete depletion of PB CM and NCM at this point. We then performed CITE-seq on CD45^+^CD11b^+^Ly6G^−^SiglecF^−^MHCII^mid/low^CD177^−^ cells 7 days post Clo-lip administration in CD45.1→CD45.2 BMC mice (Supplemental Fig 3A; Supplemental Table 3A). We labelled subpopulations using the steady state CD45.1→CD45.2 BMC CITE-seq data as reference (Fig 3C-D; Supplemental Fig 3G; Supplemental Table 3A). The composition of 3 TRMC subpopulations 7 days post Clo-lip administration was comparable to that in steady state CD45.1→CD45.2 BMC, with TIM4^+^CX3CR1^+^ TRMC and TIM4^+^ TRMC as the major subpopulations and MHCII^+^ TRMC as the minor one (Fig 3E-F). As before, we annotated the cells based on their CD45.1 and CD45.2 ADT expression (Fig 3G; Supplemental Fig 3H-I; Supplemental Table 3B). We observed a similar distribution of CD45 in each TRMC subpopulation 7 days post Clo-lip, comparable to the steady state (Fig 3H). Correlation analysis of gene expression in the three TRMC subsets, stratified by CD45 expression and excluding low cell-number populations, revealed that Day 7 post-Clo-lip TRMC subsets showed robust correlation with their corresponding steady state TRMC subsets (Supplemental Fig. 3J). Together, these results indicate that Clo-lip only primarily affected the number of host-derived TRMC while having a minimal impact on the transcriptional profiles of TRMC subpopulations 7 days post treatment.

**Figure 3.**
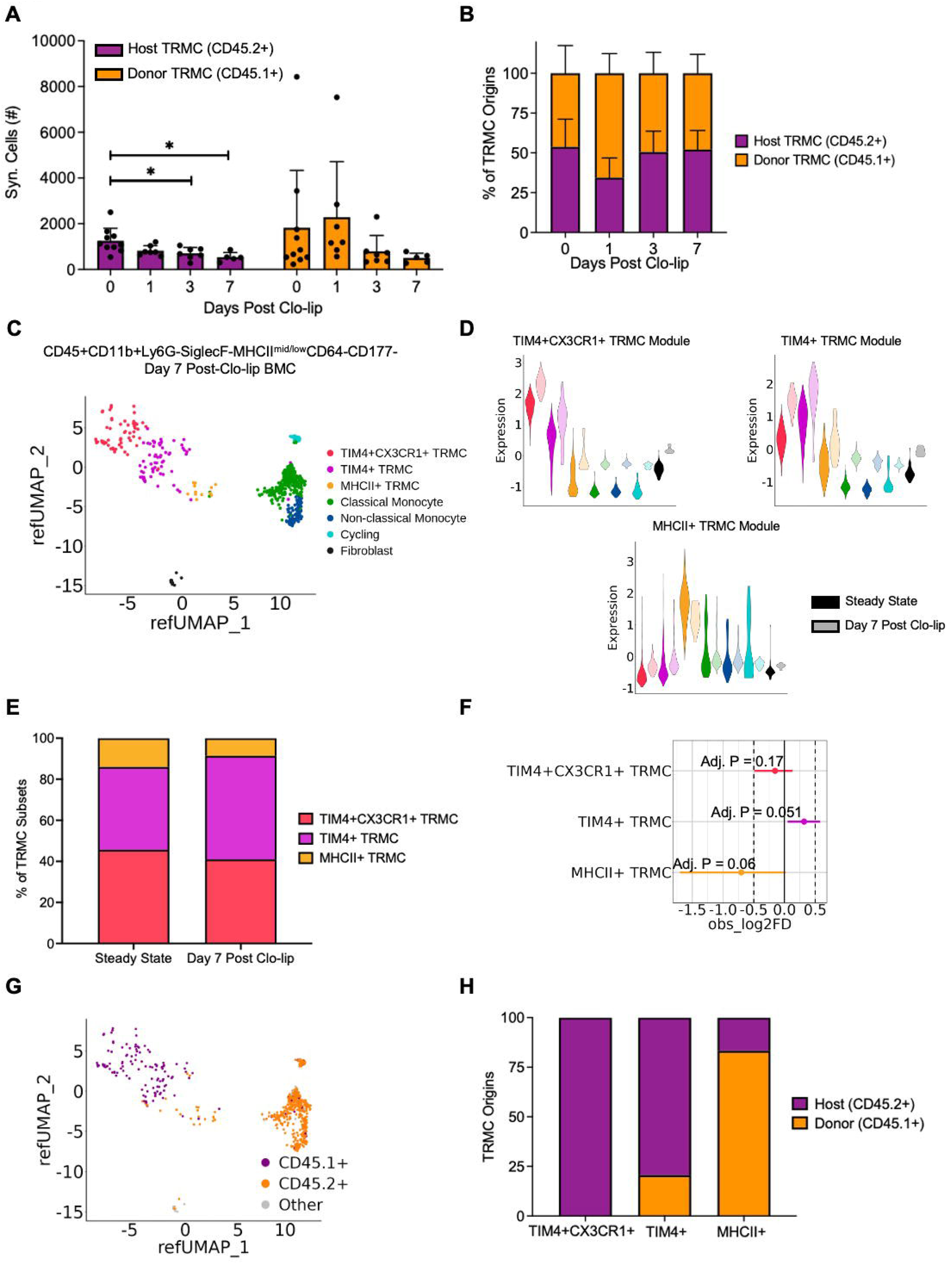
TRMC subsets from day 7 Clo-lip are transcriptionally comparable to steady state. (A) Numbers of Syn. host-derived and donor derived TRMC on day 0, 1, 3, 7 post clodronate-laden liposome (Clo-lip) administration (n ≥ 5; Treml4 was used as a proxy for I.V. CD45 to identify TRMC). Graphs were displayed as mean ± SD. P value was calculated by unpaired t-test with * p < 0.05 unless specified otherwise. (B) Percents of donor-derived and host-derived TRMC on day 0, 1, 3, 7 post Clo-lip administration (n ≥ 5). (C) Uniform manifold approximation and projection (UMAP) depicting label-transferred CD45^+^CD11b^+^Ly6G^−^SiglecF^−^MHCII^mid/low^CD64^−^ CD177^−^ cells from day 7 post Clo-lip BMC mice. (D) Violin plots of the expression of the module scores of three TRMC signature genes in steady state and day 7 post Clo-lip datasets using Fig. 2A as reference dataset. (E) Composition of 3 TRMC subsets of steady state and day 7 post Clo-lip BMC mice. (F) Comparison of the proportions of 3 TRMC subpopulations. Lines of each subpopulation represent the range of the confidence intervals from the permutation test. P values were calculated by permutation test and corrected by false discovery rate (FDR). (G) UMAP depicting CD45.1 distribution of CD45^+^CD11b^+^Ly6G^−^SiglecF^−^MHCII^mid/low^CD64^−^CD177^−^ cells from day 7 post Clo-lip BMC mice. (H) Percents of host-derived and donor-derived cells across 3 TRMC subpopulations defined in C from day 7 post Clo-lip BMC mice.

### TRMC are partially dependent on Ccr2

Since a subpopulation of TRMC (MHCII^+^ TRMC) are Ccr2^+^, we next investigated the requirement of Ccr2. As expected, PB CM were significantly depleted in Ccr2^−/−^ mice (Fig 4A). The number of TRMC was also significantly reduced in Ccr2^−/−^ mice compared to B6 (Fig 4B-C). Next, we generated reciprocal CD45.1→Ccr2^−/−^ and Ccr2^−/−^ →CD45.1 BMC mice to assess whether the absence of Ccr2 affected the numbers of donor-derived or host-derived TRMC (Fig 4D). Analysis of PB showed >95% of chimerism in PB CD45^+^, PB CM, and PB NCM (Supplemental Fig 4A), along with a significant depletion of both PB CM and PB NCM in CD45.1→Ccr2^−/−^ compared to Ccr2^−/−^ →CD45.1 BMC mice, due to the absence of Ccr2 in the bone marrow (Supplemental Fig 4B). The total number of TRMC remained comparable between Ccr2^−/−^ →CD45.1 and CD45.1→Ccr2^−/−^ BMC mice (Fig 4E). However, compared to steady state, the ratio of donor-derived to host-derived TRMC was significantly reduced, in Ccr2^−/−^ →CD45.1 BMC mice (Fig 4E; Supplemental Fig 4C). On the other hand, CD45.1→Ccr2^−/−^ BMC mice exhibited similar TRMC numbers to steady-state CD45.1→CD45.2 BMC mice (Fig 4E). Notably, while CD45.1→Ccr2^−/−^ BMC mice exhibited comparable surface marker expression of TIM4, CX3CR1, and MHCII to Ccr2^−/−^ →CD45.1 BMC, host-derived TRMC from Ccr2^−/−^ →CD45.1 exhibited a lower proportion of TIM4^+^ cells and a proportion of higher MHCII^+^ cells (Supplemental Fig 4D). However, the numbers of MHCII^+^ TRMC were comparable to Ccr2^−/−^ →CD45.1 BMC mice when combined with donor cells (Supplemental Fig 4E), suggesting the expansion of host-derived TRMC are compensating for the reduced donor-derived MHCII^+^ TRMC. These findings suggest that donor-derived TRMC, but not host-derived, are dependent on Ccr2.

**Figure 4.**
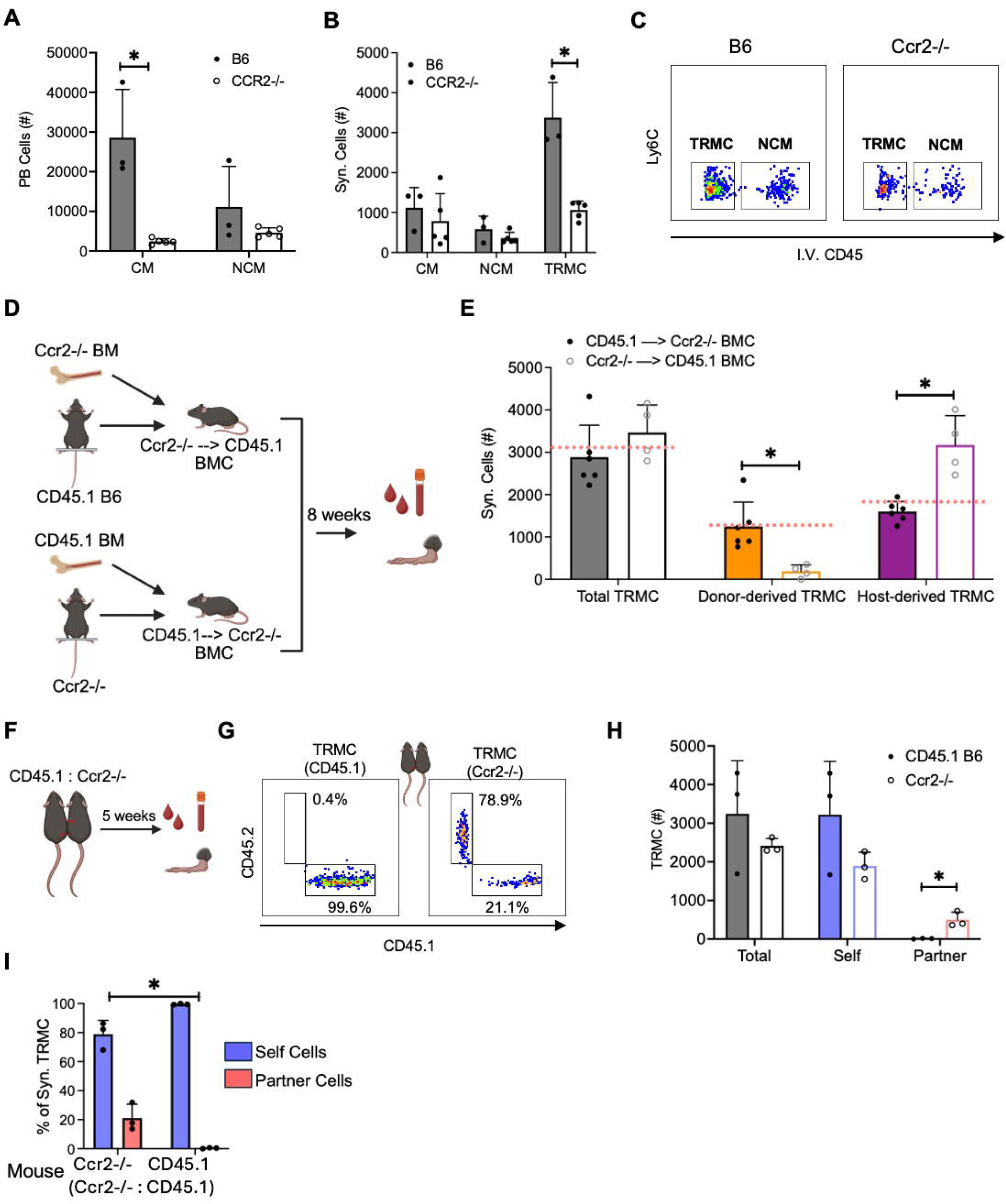
Donor-derived TRMC is Ccr2-dependent. (A) Numbers of PB CM and NCM in B6 and Ccr2^−/−^ mice (numbers are representatives of the averages of n ≥ 3; (B) Numbers of Syn. CM, NCM, and TRMC in B6 and Ccr2^−/−^ mice (numbers are representatives of the averages of n ≥ 3; Treml4 was used as a proxy for I.V. CD45 to identify TRMC). (C) The distribution of TRMC and NCM in B6 and Ccr2^−/−^ mice. (D) Schematic of reciprocal Ccr2^−/−^→CD45.1 and CD45.1→Ccr2^−/−^ BMC mice. (E) Numbers of total, donor-derived, and host-derived TRMC in Ccr2^−/−^→CD45.1 (n = 4) and CD45.1→Ccr2^−/−^ BMC (n = 6). Red dash lines representing the average numbers of TRMC from steady state CD45.1→CD45.2 BMC shown in Fig. 3. (F) Schematic of CD45.1:Ccr2^−/−^ parabiosis mice. (G) Percents of CD45.1^+^ and CD45.2^+^ TRMC in CD45.1:Ccr2^−/−^ parabiosis mice (numbers are representatives of the averages of n ≥ 3; Treml4 was used as a proxy for I.V. CD45 to identify TRMC). (H) Numbers of total, self, partner TRMC in CD45.1:Ccr2^−/−^ parabiosis mice (n = 3). (I) Percents of self and partner cells of Syn. CM, NCM, and TRMC in CD45.1:Ccr2^−/−^ parabiosis mice (n = 3). Chi-square test was used to compare the distribution of cell origins from Syn. tissue between CD45.1 B6 and Ccr2^−/−^ from CD45.1:Ccr2^−/−^ parabiosis, under the null hypothesis that the distribution of cell origins of any cell type does not different between the two mice groups in Syn. tissue. All graphs are displayed as mean ± SD. P value was calculated by unpaired t-test with * p < 0.05.

Next, we generated CD45.1:Ccr2^−/−^ parabiotic mice (Fig 4F; Supplemental Fig 4F). As expected, the number of PB CM was lower in the Ccr2^−/−^ parabiont with a higher proportion of partner-derived cells (Supplemental Fig 4G). The percentage of partner-derived TRMC was markedly higher in Ccr2^−/−^ parabiont (21.1%) of CD45.1:Ccr2^−/−^ parabiosis compared to CD45.2 parabiont of CD45.1:CD45.2 parabiosis (3.7%) (Fig 1F-G; Fig 4G). Moreover, the number of partner-derived TRMC was significantly higher in Ccr2^−/−^ parabiont compared to CD45.1 parabiont, while the total and self-derived TRMC remained comparable (Fig 4H). The distribution of PB monocytes and TRMC showed a significant difference between Ccr2^−/−^ and CD45.1 parabionts (Fig 4I; Supplemental Fig 4H). These data suggest that the bone marrow-derived TRMC are replenished by partner-derived cells when the self-bone marrow lacks Ccr2, further confirming the survival dependent of a TRMC subpopulation on Ccr2.

### Donor-derived TRMC expanded during the development of STIA

To investigate the impact of inflammatory arthritis on TRMC composition, we isolated TRMC from different stages of STIA. Numbers of host-derived and donor-derived PB CM, PB NCM, Syn. CM, Syn. NCM, and TRMC were measured in CD45.1→CD45.2 BMC mice during the baseline (day 0), peak (day 11), plateau (day 16), and resolution (day 56) stages of STIA (Supplemental Fig 5A-B). CM and NCM from PB and synovial tissue expanded during STIA progression except donor-derived PB NCM with a reduction on peak and plateau compared to baseline (Supplemental Fig 5C-F). This reduction may reflect enhanced recruitment of PB NCM to inflamed synovial tissue. Host-derived TRMC were significantly depleted during the peak, plateau, and resolution stages of STIA, whereas donor-derived TRMC were significantly expanded, showing more than a two-fold increase in number at both the peak and plateau stages (Fig 5A-B). By the resolution stage, the numbers of both host-derived and donor-derived TRMC exhibited a dynamic return toward levels observed at the steady state (Fig 5A-B). While over half of TRMC were host-derived at steady state, the majority shifted to donor-derived during peak and plateau stages (Fig 5C). This ratio gradually returned to the steady state composition by the resolution stage (Fig 5C). TIM4 and MHCII expression levels were markedly lower at STIA peak and plateau in host-derived and donor-derived TRMC, respectively (Fig 5D). In contrast, CX3CR1 expression did not differ between the two TRMC subsets during the peak, plateau, or resolution stages (Fig 5D). Quantification of TIM4⁺, CX3CR1⁺, and MHCII⁺ TRMC revealed that donor-derived MHCII⁺ TRMC were the major contributors during the peak and plateau stages of STIA, accompanied by a reduction in host-derived TIM4⁺ and CX3CR1⁺ TRMC (Fig 5E). Collectively, these findings indicate that synovial inflammation leads to significant expansion of donor-derived TRMC and depletion of host-derived TRMC, accompanied by phenotypic alterations.

**Figure 5.**
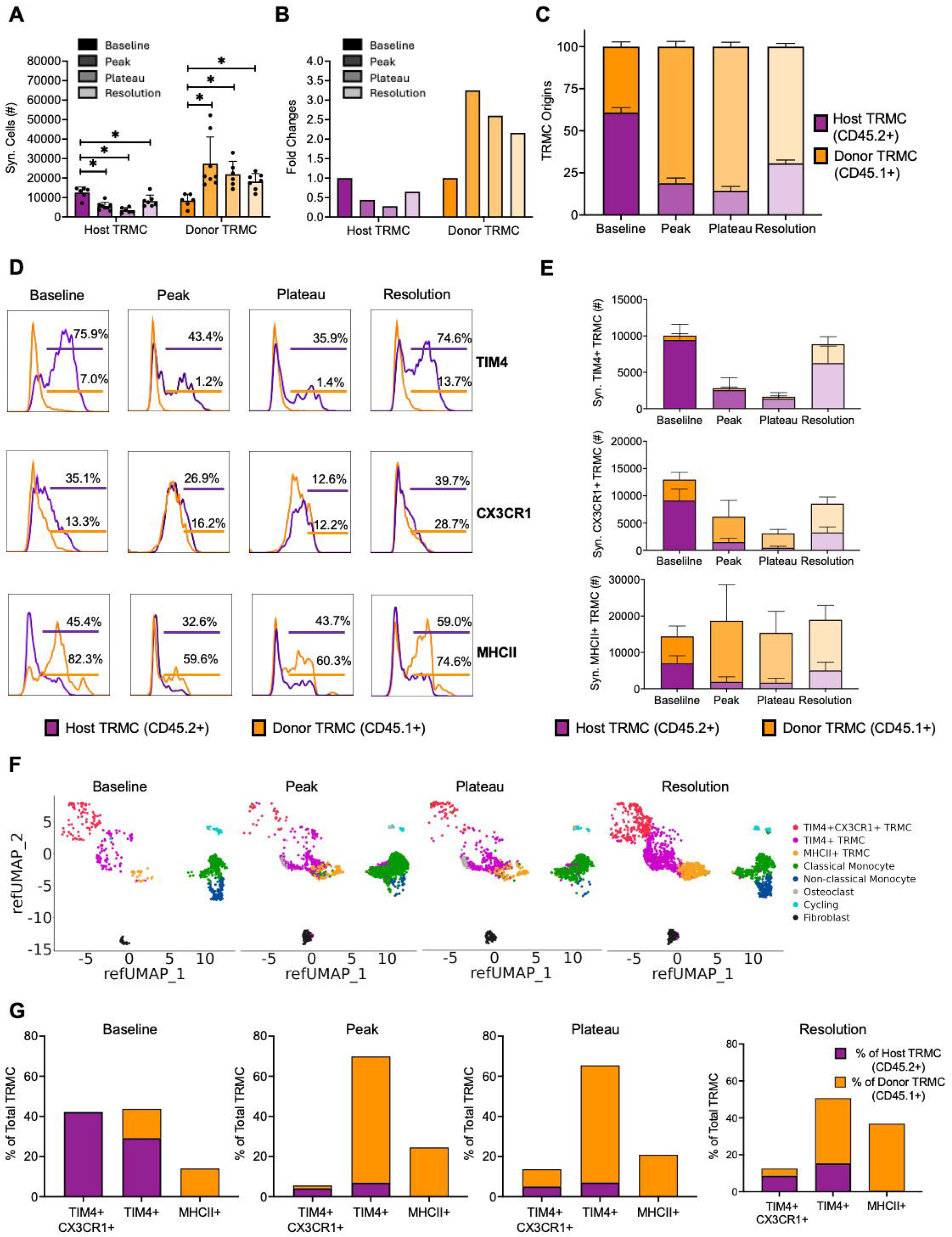
The numbers and proportions of TRMC subpopulations undergo dynamic changes during the development of STIA. (A) Numbers of Syn. host-derived and donor derived NCM from the baseline, peak, plateau, and resolution stages of K/BXN serum-transfer-induced arthritis (STIA) development (n ≥ 5). (B) Fold changes of Syn. host-derived and donor derived TRMC from the baseline, peak, plateau, and resolution stages of STIA development (n ≥ 5). (C) Percents of donor-derived and host-derived TRMC from the baseline, peak, plateau, and resolution stages of STIA development (n ≥ 5). (D) Expression of TIM4, CX3CR1, and MHCII of host-derived and donor-derived TRMC from the baseline, peak, plateau, and resolution stages of STIA development. (E) Numbers of TIM4^+^, CX3CR1^+^, MHCII^+^ TRMC split by CD45 expression from the baseline, peak, plateau, and resolution stages of STIA development, respectively. (F) Annotation of synovial cell populations of CD45^+^CD11b^+^Ly6G^−^SiglecF^−^MHCII^mid/low^CD64^−^CD177^−^ cells of BMC mice from the baseline, peak, plateau, and resolution stages of STIA development, respectively. (G) Proportions of origins of three TRMC subpopulations in the four STIA stages, respectively. All graphs were displayed as mean ± SD. P value was calculated by unpaired t-test with * p < 0.05.

### STIA-associated TRMC exhibit diverse transcriptional phenotypes

We performed CITE-seq on CD45^+^CD11b^+^Ly6G^−^SiglecF^−^MHCII^mid/low^CD177^−^ cells from CD45.1→CD45.2 BMC mice during the development of STIA (Supplemental Fig 5A). Using the steady state CITE-seq dataset annotated in Fig 2A as reference, we identified seven cell types across four STIA datasets. However, there was significant expression of osteoclast-associated genes during STIA, as noted previously ^6^. Therefore, we independently annotated these cells (Supplemental Fig 5G) and determined that they were found almost exclusively in the peak and plateau stages of STIA (Fig 5F).

Next, we investigated how the distribution of CD45.2^+^ (host-derived) and CD45.1^+^ (donor-derived) cells varied within TRMC subpopulations throughout STIA progression (Supplemental Fig 5H-I; Supplemental Table 4A-D). As expected, synovial CM and NCM were exclusively donor-derived at all stages of STIA (Supplemental Fig 5I). Consistent with the reference dataset, host-derived TRMC represented the predominant subset at baseline within total TRMC, TIM4^+^CX3CR1^+^ TRMC, and TIM4^+^ TRMC, while MHCII^+^ TRMC were exclusively donor-derived (Fig 5G). In the peak stage of STIA, donor-derived TRMC became the predominant population among total TRMC, although host-derived TRMC remained the major subset in TIM4^+^CX3CR1^+^ TRMC (Fig 5G). During the plateau stage of STIA, donor-derived TRMC became the dominant population across all three subsets (Fig 5G). With resolution, host-derived proportions of total, TIM4^+^CX3CR1^+^, and TIM4^+^ TRMC rebounded but did not quite reach baseline levels (Fig 5G). The changes observed in the numbers and distribution of host-derived and donor-derived TRMC throughout STIA progression were consistent between the flow cytometry and CITE-seq datasets. Together, these results demonstrate that synovial inflammation led to the recruitment of donor-derived TRMC and a decrease in tissue-resident phenotype.

### Synovial inflammation alters transcriptional profiles of TRMC subsets

To identify functional changes in TRMC across the course of STIA, we compared their transcriptional profiles across timepoints. We found that TRMC subsets from the baseline and resolution stage of STIA exhibited comparable expression of subset-specific genes to those in the reference dataset (Supplemental Fig 6A). However, the peak and plateau stages of STIA were associated with a loss of these steady-state signatures (Fig 6A; Supplemental Fig 6A). Donor-derived TIM4⁺ TRMC from the peak and plateau stages lacked Tim4 (ADT) expression, whereas host-derived TRMC retained it. Both donor- and host-derived TIM4⁺ TRMC exhibited low levels of I-A-I-E (MHCII) expression as the baseline stage. Host-derived (CD45.2^+^) and donor-derived (CD45.1^+^) TIM4^+^ TRMC from the peak and plateau stages were somewhat divergent but exhibited a distinct transcriptional signature from other subpopulations (Figure 6B). When we clustered 1000 most variable genes based on their pseudo bulk expression across subsets and time points, we identified 5 distinct transcriptional trends (Figure 6C; Supplemental Table 5). Cluster 1 showed the highest expression in TIM4^+^CX3CR1^+^ TRMC across all datasets and in TIM4^+^ TRMC from the baseline and resolution stages (Figure 6C). Cluster 1 was associated with complement activation pathway, featuring genes such as C1qa, C1qb, and C1qc, as well as TIM4^+^CX3CR1^+^ TRMC specific DEG Vsig4 and TIM4^+^ TRMC specific DEG Aqp1 in the steady state BMC dataset (Figure 6C; Supplemental Fig 2B; Supplemental Fig 6B; Supplemental Table 6A). Cluster 2 genes were specific to MHCII^+^ TRMC across all 4 datasets and in TIM4^+^ TRMC from the peak and plateau stages, featuring genes (Adam8, Ccr2) were associated with pathways such as leukocyte chemotaxis (Fig. 6C; Supplemental Fig 6B; Supplemental Table 6B). Genes, including Cd74, H2-Aa, and Cd24a (Supplemental Fig 6B), showed the highest expression in Cluster 3 which was significantly associated with antigen processing and presentation of exogeneous antigen pathway and enriched in TIM4^+^ TRMC from the peak stage and in MHCII^+^ TRMC except the plateau stage (Fig. 6C; Supplemental Table 6C). Cluster 4 (Apoe, Hif1a, Arg1) and cluster 5 (Cd81, Mef2c, Alox5) represented signature genes of the peak and plateau stages, and the baseline and resolution stages, respectively (Fig 6C; Supplemental Fig 6B). Cluster 4 contained genes associated with wound healing, and Cluster 5 genes were associated myeloid leukocyte differentiation (Fig 6C; Supplemental Fig 6B; Supplemental Table 6E). Associated pathways were generally validated using a single-cell GSVA approach, except in the peak and plateau stages, where the minority CD45.2^+^ TIM4^+^ TRMC exhibited a similar enrichment pattern to TIM4^+^CX3CR1^+^ (Supplemental Fig. 6C; Supplemental Table 7).

**Figure 6.**
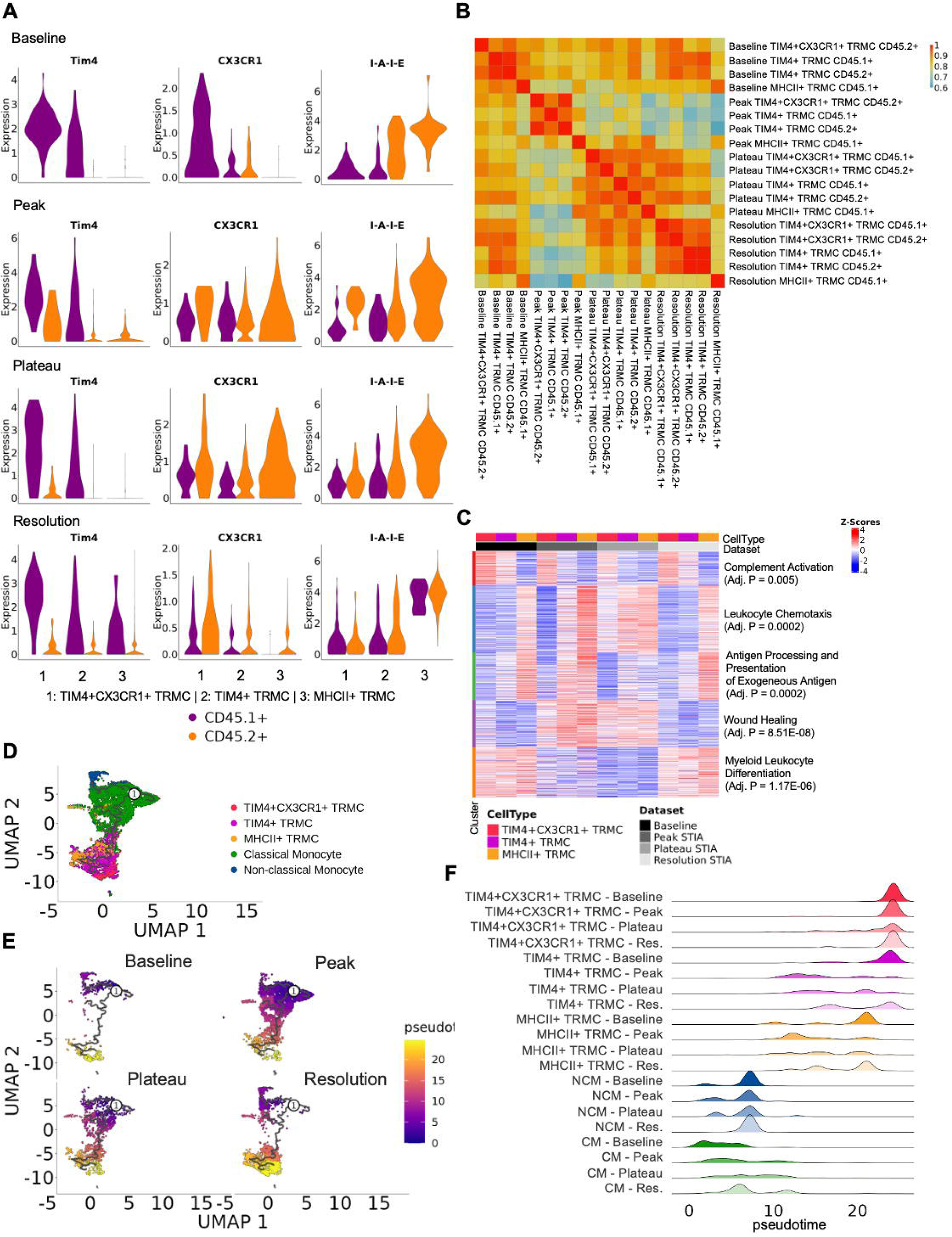
STIA development alters the functions of TRMC subsets. (A) Violin plots of the ADT expression of Tim4, CX3CR1, I-A-I-E (MHCII) of the three TRMC subsets stratified by their CD45 expression from the four stages of STIA development. (B) Heatmap showing the correlation of the three TRMC subsets stratified by CD45 expression across the four stages of STIA development. (C) Heatmap of K-means clustering results of pseudo bulk analysis of the three TRMC subsets across the four stages of STIA development. (D) UMAP depicting the annotation of monocyte-lineage cell types (CM, NCM, and three TRMC subsets) across the four stages of STIA development. (E) UMAP depicting the pseudo time trajectory of merged monocyte-linage cells across the four stages of STIA development. (F) Ridge plot of the pseudo time values of the three TRMC subsets across the four stages of STIA development.

Finally, we performed trajectory analysis on monocyte populations, including CM, NCM, and all TRMC subsets across the STIA datasets. We defined CM as the starting point, given that NCM are derived from CM ^28^. Our results were consistent with a branching developmental path originating from CM: one branch progressed toward NCM, while the other diverged into TIM4^+^ TRMC and MHCII^+^ TRMC. Subsequently, TIM4^+^ TRMC moved into TIM4^+^CX3CR1^+^ TRMC (Fig 6D-E). The stage of STIA affected the pseudotime values of TRMC: while baseline TIM4^+^ TRMC and MHCII^+^ TRMC exhibited a single peak, these subpopulations exhibited a range of lower values during inflammation suggesting they exist at intermediate states. By resolution, the majority of these populations have returned to their baseline pseudotime values, but an intermediate subpopulation still persisted (Fig 6F). These findings suggest that the functions of the TRMC subset are altered following the development of STIA with the majority donor-derived TIM4^+^ TRMC exhibiting inflammatory pathways and features of MHCII^+^ TRMC that are downregulated following resolution.

## Discussion

Our study investigated the heterogeneity and ontogeny of TRMC, a monocyte-lineage cell type residing in the synovial tissue. We found that TRMC are composed of three distinct subsets: TIM4^+^CX3CR1^+^, TIM4^+^, and MHCII^+^. TIM4^+^CX3CR1^+^ and TIM4^+^ are primarily long-lived and MHCII^+^ are bone-marrow (BM)-derived. Following Clo-lip administration, TIM4^+^CX3CR1^+^ TRMC and TIM4^+^ TRMC are predominantly long-lived cells, whereas BM-derived cells replenish their niches during STIA development. Regardless of the inflammatory states, MHCII^+^ TRMC are entirely BM-derived cells and dependent on Ccr2. Furthermore, we demonstrated that TRMC subsets exhibit distinct transcriptional profiles, surface expression of TIM4, CX3CR1, and MHCII, as well as functional differences that vary with treatment and inflammatory context. Together, our findings suggest that TRMC consist of three transcriptionally and functionally distinct subsets that play unique roles in the pathogenesis of inflammatory arthritis.

To investigate the origins of TRMC, we used BMC and parabiosis mice. During bone marrow chimera generation, shielded irradiation is typically used to protect the tissue resident cells. We previously used CD45.1→CD45.2 BMC mice, with ankles shielded to preserve the tissue-resident cells, to demonstrate that MHCII^+^ synovial macrophages are BM-derived ^7^. Similarly, by using reciprocal CX3CR1^CreER^.zsGFP→C57BL/6 (B6) and B6→CX3CR1^CreER^.zsGFP, with tamoxifen injected on embryonic day 15, TRMC was proven to be at least partially embryonic-derive ^6^. Parabiotic mice are another useful method for studying the origins of cells. For example, Ginhoux F, *et al.,* used parabiosis mice to show that adult microglia are maintained ^29^. In synovial tissue, Culemann S, *et al*. used CD45.1 B6:Cx3cr1^GFP^ to prove that CX3CR1^+^ lining macrophages are independent of circulating monocytes ^8^. Together, BMC and parabiosis models serve as valuable tools for studying the origins of specific cell types and have been used in this study to demonstrate the proportions of BM-derived vs. long-lived TRMC.

Mononuclear phagocytes, specifically macrophages, are a group of cells with a high level of heterogeneity in both human and mouse that was studied extensively using methods such as single-cell RNA sequencing (scRNA-seq)^7–10,29–41^. For example, Kupffer cells, as one of the earliest discovered tissue-resident yolk-sac derived macrophages ^35^, have recently been reported to have two subsets distinguished by the expression of Marco using scRNA-seq, with the Marco⁺ subset enriched in the periportal vein zone of the liver and exhibiting anti-inflammatory properties ^30^. Specifically in synovial tissue, Culemann S, *et al.* identified CX3CR1^+^ tissue-resident macrophages, which are enriched in the synovial lining and have been reported to form a protective barrier against inflammatory arthritis, along with three additional interstitial macrophages identified using scRNA-seq ^8^. Subpopulations of synovial macrophages with distinct contributions to rheumatoid arthritis (RA) development have also been identified in human tissues ^8,33,34^, such as MerTK^pos^TREM2^pos^ synovial tissue macrophages, which are homologous to murine CX3CR1^+^ lining macrophages and are found in the synovium of healthy donor and RA patients in remission ^33^. Additionally, we previously showed the homologous compartment of TRMC are CD68^+^TREM2^−^TIM4^+^ synovial cells in healthy human tissue, which becomes CD163^+^ in RA patients ^6^. These findings highlight the heterogeneity of mononuclear phagocytes across different tissues and underpin the hypothesis that TRMC as a synovial tissue resident mononuclear phagocyte population that are comprised of different subsets. Therefore, we performed CITE-seq on CD45+CD11b+Ly6G-SiglecF-MHCII^mid/low^CD64-CD177-synovial cells. Annotation of the steady state BMC dataset using transcriptional and surface markers enabled us to identify three TRMC subsets, distinguished by the surface expression of TIM4, CX3CR1, and MHCII. These findings were further validated with flow cytometry results.

CM rely on Ccr2, the receptor for monocyte chemoattractant protein-1 (MCP-1), for their exit from the bone marrow and recruitment to inflamed tissues ^42,43^. Depletion of Ccr2 significantly reduces PB CM and I-NCM in lung, however, its effect on inflammatory arthritis in mice has been reported with conflicting outcomes, depending on the specific mouse models ^6,7,14,44,45^. It has no effect in the K/BxN serum-transfer-induced arthritis (STIA) model ^6^ but significantly exacerbates inflammation in the human TNF-transgenic (hTNFtg) ^44,46^. Ccr2 deficiency has been shown to reduce the number of Ccr2+ macrophages compared with wild type (WT) in peripheral nervous system ^31^, MHCII^+^ ^47^, F4/80^+^ ^48^, F4/80^low^ liver ^49^, and F4/80^+^CD11b^+^ ^49^, presumably because these macrophages are monocyte-derived ^50,51^. In our studies, we showed that the BM-derived TRMC are CCR2-dependent, and this is the subpopulation driving TRMC expansion in STIA. These findings underscore the critical role of Ccr2 in monocyte survival and the regulation of monocyte-derived cell populations, which may contribute to the varying effects of Ccr2 deficiency across different inflammatory arthritis models.

Previously, we have reported that TRMC were not dependent on Ccr2. However, in that study, we only calculated the impact on total TRMC numbers. Moreover, further investigation later revealed that the previously identified TRMC population may have included Ly6C^low^CD177^+^ population, which are believed to be Ly6G^−^ neutrophils, potentially residual neutrophils from the BM ^16,52^. CITE-seq analysis of the steady state BMC dataset in this study revealed that MHCII^+^ TRMC express Ccr2, prompting us to use Ccr2^−/−^ mice to investigate whether the maintenance of TRMC subpopulations depends on Ccr2. Utilizing Ccr2^−/−^ and CD45.1 reciprocal BMC mice, we showed that Ccr2 deficiency greatly reduced the donor-derived TRMC subpopulations in Ccr2^−/−^→ CD45.1 BMC compared to CD45.1→Ccr2^−/−^ or CD45.1→CD45.2 BMC. This reduction suggests that donor-derived TRMC are Ccr2-dependent. Interestingly, we also observed reduced TIM4 and increased MHCII expression in host-derived TRMC when donors lacked Ccr2. This discrepancy suggests a compensatory mechanism, in which the newly expanded host-derived TRMC phenotypically resemble the absent donor-derived TRMC in the context of Ccr2 deficiency. This may explain why Ccr2^−/−^ mice are able to develop arthritis despite our prior results suggesting that an expanded TRMC population is necessary for STIA development ^6^.

The progression of inflammation can induce the expression of previously unexpressed genes, thereby altering the transcriptional profile of monocytes/macrophages. For example, Culemann S, et al. identified synovial inflammation-specific Ccr2^+^ infiltrating macrophages ^8^, which are homologous to human IL1B^+^ proinflammatory monocytes that enriched in lipopolysaccharide (LPS) stimulation-related pathways ^34^. Human MerTK^pos^FOLR2^+^ID2^+^ synovial tissue macrophages (STM) exhibited enrichment in complement & defensin system pathways in healthy controls, but become transcription-factor activation pathway-enriched in RA patients ^33^. Similarly, we observed that TIM4^+^ TRMC lost their characteristic transcriptional signature during the peak and plateau stages of STIA and gained MHCII^+^ TRMC signatures, which was corroborated by both GSVA results and pseudo bulk RNA-seq analysis. Pseudo time analysis of the monocyte subpopulations during STIA suggested that TIM4^+^ TRMCs exist in an intermediate stage between circulating monocytes and their baseline state at the peak and plateau inflammation. Collectively, these suggest that the onset, progression, and resolution of inflammation in synovial tissue alters the phenotype of TRMC.

One missing piece of the puzzle in this study is the lack of a specific depletion model to assess the role of TRMC in inflammatory arthritis. TRMC. While TRMC, particularly those long lived, are sensitive to Clo-lip, batch effects altering the efficacy between trials are also evident ^6^. GO results of three TRMC subsets revealed distinct functional profiles both at steady state and during inflammation, underscoring the critical roles they play in synovial tissue. We screened 18 available myeloid-related lineage-tracing mouse models and found that TRMC were labeled at varying efficiencies, with around either 10-40 % or 60-90% in each model, suggesting that different subsets of TRMC are targeted by these models ^16^. Additionally, we have identified several potential specific TRMC-specific markers and are currently in the process of generating transgenic mouse lines to enable selective depletion ^16^. Future experiments will incorporate appropriate depletion strategies to selectively eliminate all or specific TRMC subsets, thereby allowing for dissection of their functional contributions to STIA development.

## Acknowledgments

We would like to thank the Northwestern University Lurie Cancer Center Flow Cytometry Core Facility, supported by NCI Cancer Center Support Grant P30 CA060553. We would also like to thank Northwestern University Microsurgery and Preclinical Research Core Facility. This research was supported in part through the computational resources and staff contributions provided for the Quest high performance computing facility at Northwestern University and computational resources and staff contributions provided by the Genomics Compute Cluster which are both jointly supported by the Feinberg School of Medicine, the Center for Genetic Medicine, and Feinberg’s Department of Biochemistry and Molecular Genetics, the Office of the Provost, the Office for Research, and Northwestern Information Technology. The Genomics Compute Cluster is part of Quest, Northwestern University’s high performance computing facility, with the purpose to advance research in genomics. The study was supported by the NIH through R01 AI163742 (Deborah R. Winter), R21 AR080351 (Deborah R. Winter), R01 AR080513 (Harris Perlman & Deborah Winter), and R01 AR075423 (Harris Perlman) as well as by the Hevolution/AFAR New Investigator Award (DRW).

## Materials and Software

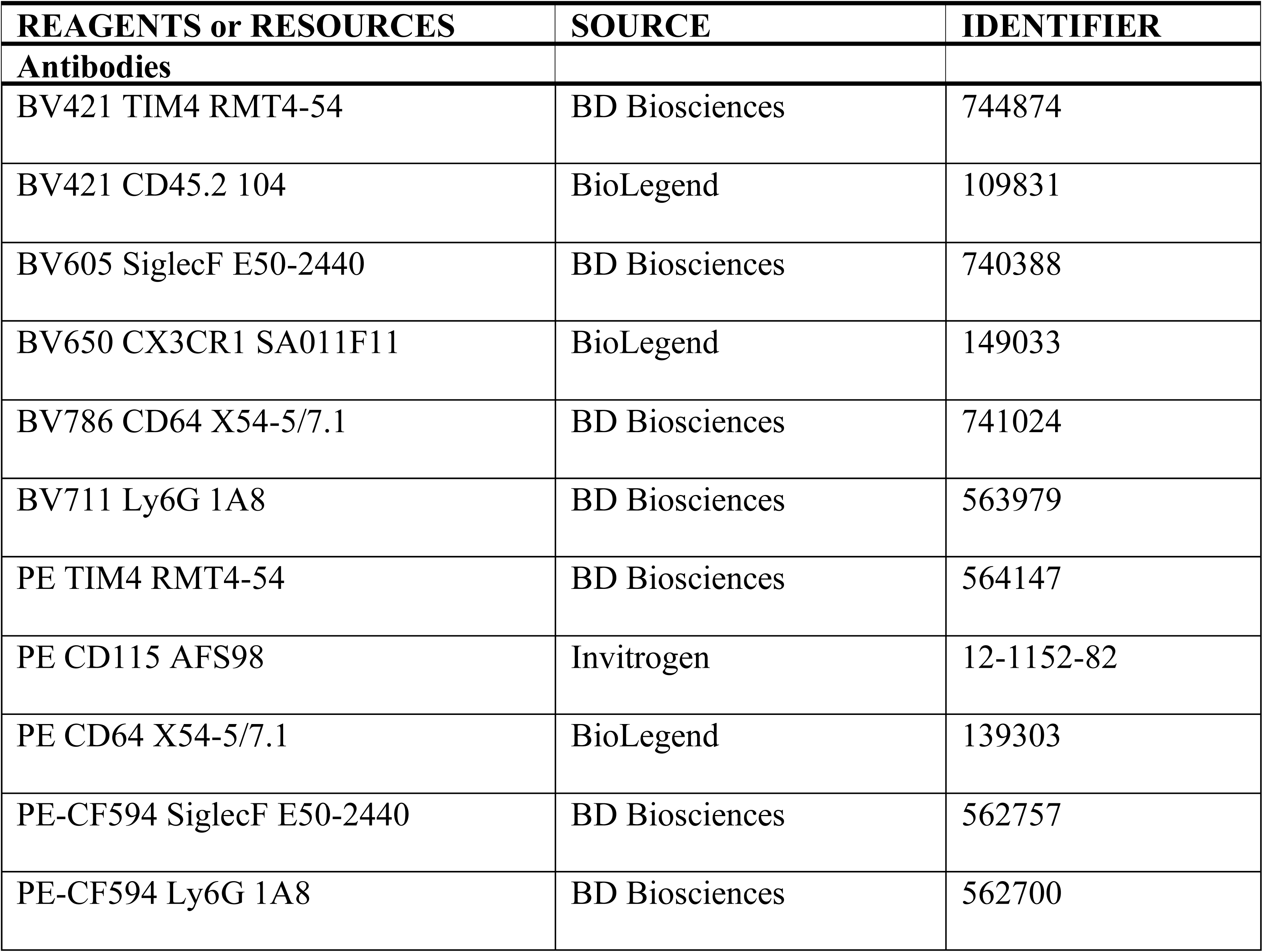
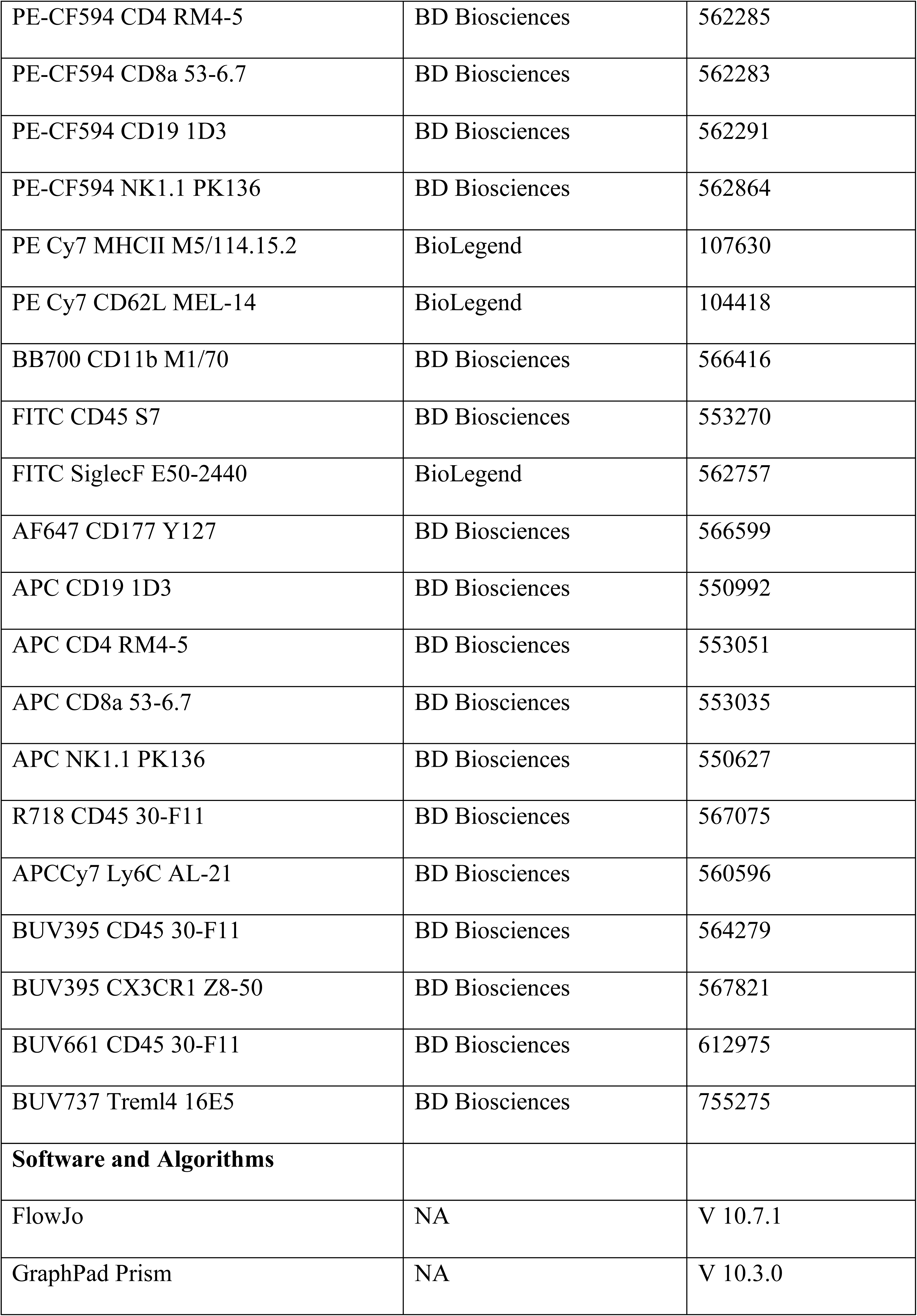
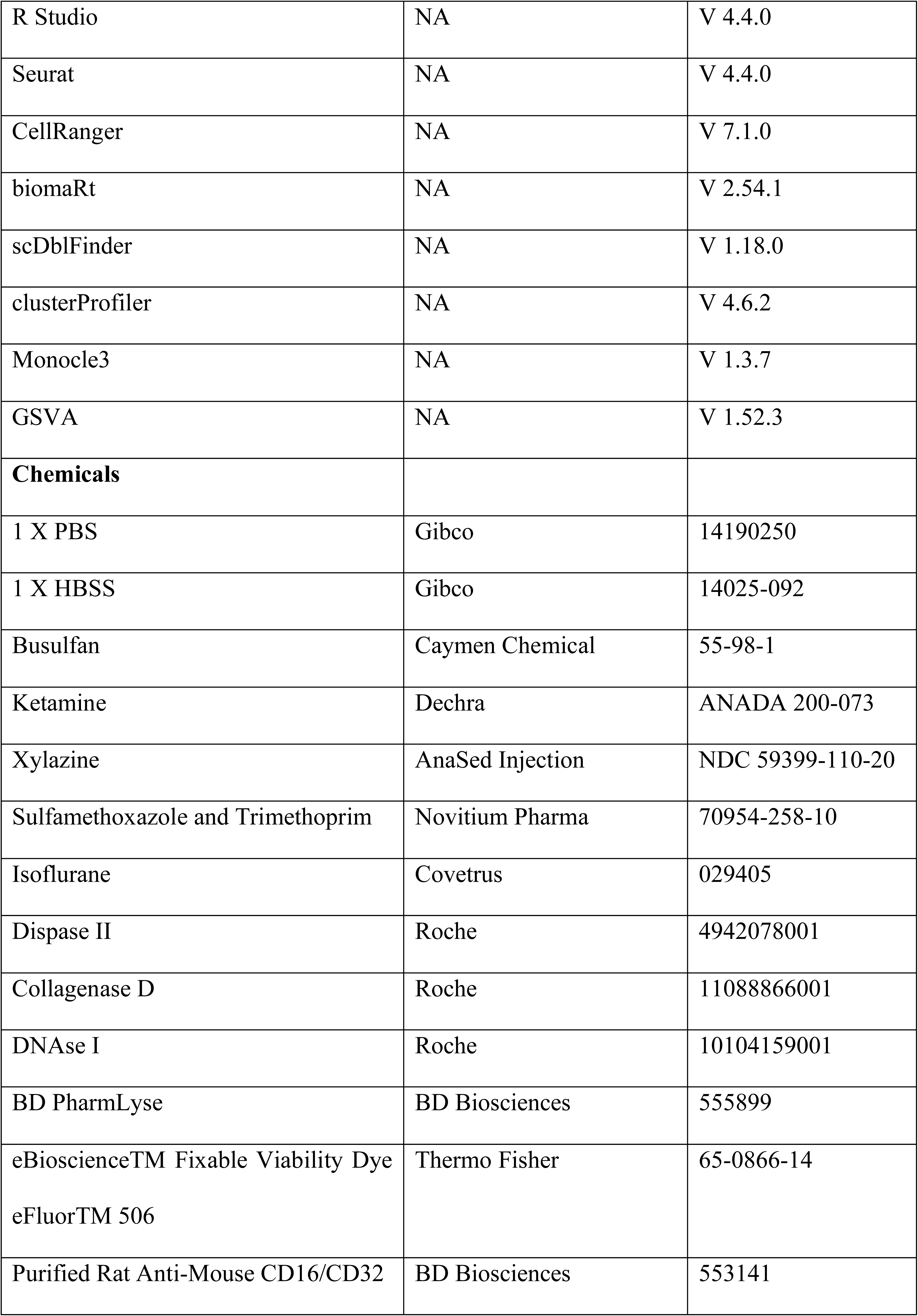
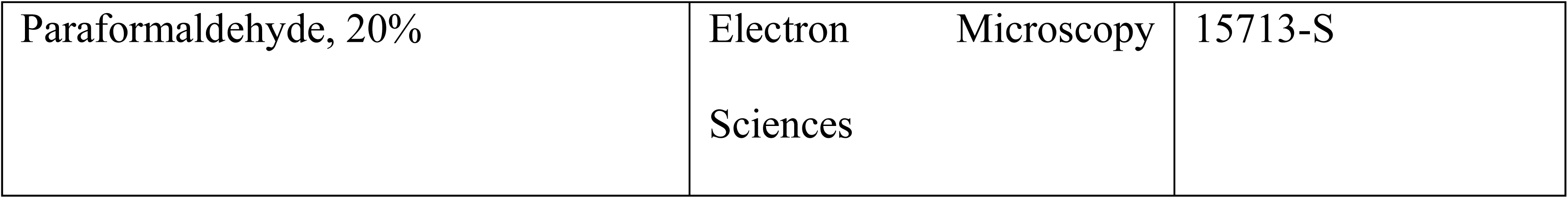

---

**Supplemental Figure 1.**
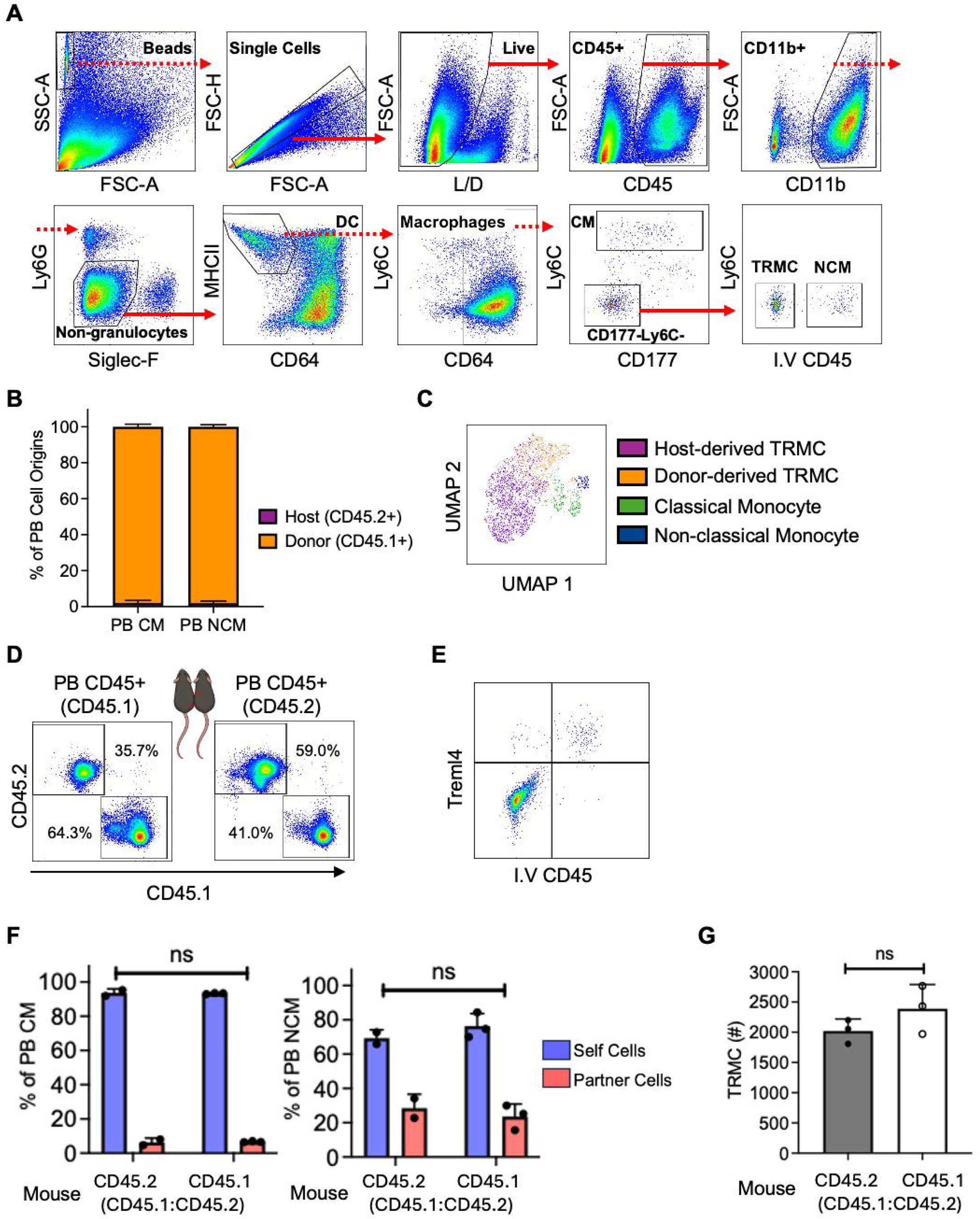
Identification of TRMC using flow cytometry. (A) Gating strategy for the identification of TRMC in CD45.1→CD45.2 BMC synovial tissue. Solid arrows indicate positive gating, while dashed arrows indicate negative gating. (B) Percent of host-derived and donor-derived PB CM and PB NCM in steady state CD45.1→CD45.2 BMC (n= 27). (C) Uniform Manifold Approximation and Projection (UMAP) depicting the distribution of host-derived TRMC, donor-derived TRMC, classical monocytes, and non-classical monocyte based on all the surface markers excluding CD45.1 and CD45.2. (D) Expression of CD45.1 and CD45.2 of PB CD45^+^ from CD45.2 B6 and CD45.1 B6 in CD45.1:CD45.2 parabiosis mice (numbers are representatives of the averages of n ≥ 2). (E) Expression of I.V CD45 and Treml4 for Syn. TRMC. (F) Percents of CD45.1^+^ and CD45.2^+^ cells of PB CM and NCM in CD45.1:CD45.2 B6 parabiosis mice (n = 3). Ns indicates p-value > 0.05 (chi-square test). (G) Number of total TRMC from CD45.2 B6 and CD45.1 B6 in CD45.1:CD45.2 parabiosis mice (n = 3). Ns indicates p-value > 0.05 (t test).

**Supplemental Figure 2.**
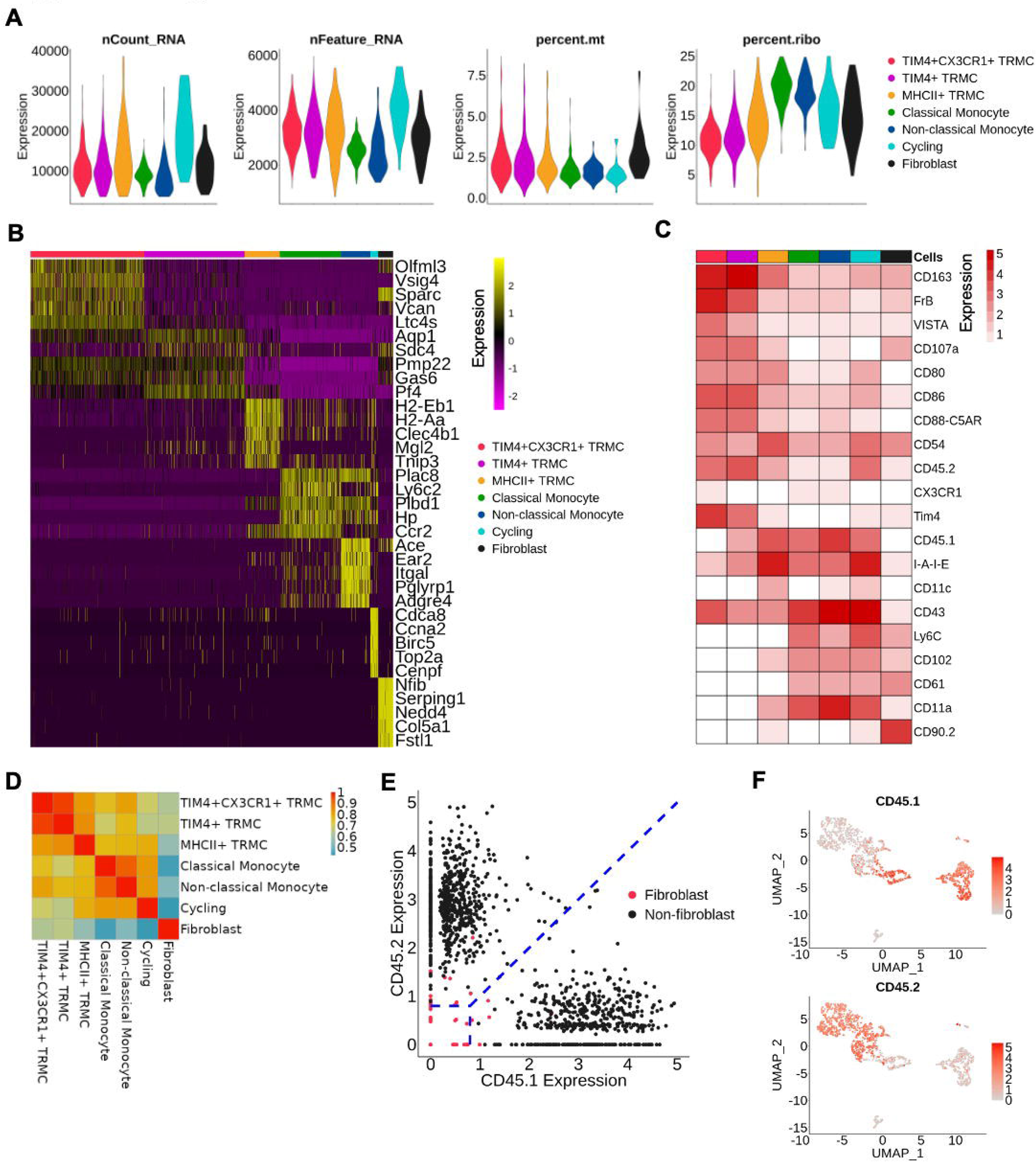
Three TRMC subpopulations display distinct transcriptional and ADT profiles. (A) Violin plots of nCount_RNA, nFeature_RNA, percent.mt, and percent.ribo of seven annotated populations identified in Figure. 2A. (B) Heatmap of the top 5 differentially expressed genes (DEG) for seven annotated populations identified in Figure. 2A. (C) The average expression of antibody-derived tags (ADT) surface markers across the seven annotated populations identified in Figure. 2A. (D) Pairwise Pearson correlation plot showing the average expression across the seven annotated populations identified in Figure. 2A. (E) The classification of CD45.1^+^ and CD45.2+ cells for fibroblast and non-fibroblast cells from CD45^+^CD11b^+^Ly6G^−^SiglecF^−^ MHCII^−^CD64^−^CD177^−^ cells. (F) Feature plots of the expression of CD45.1 and CD45.2 (ADT surface marker).

**Supplemental Figure 3.**
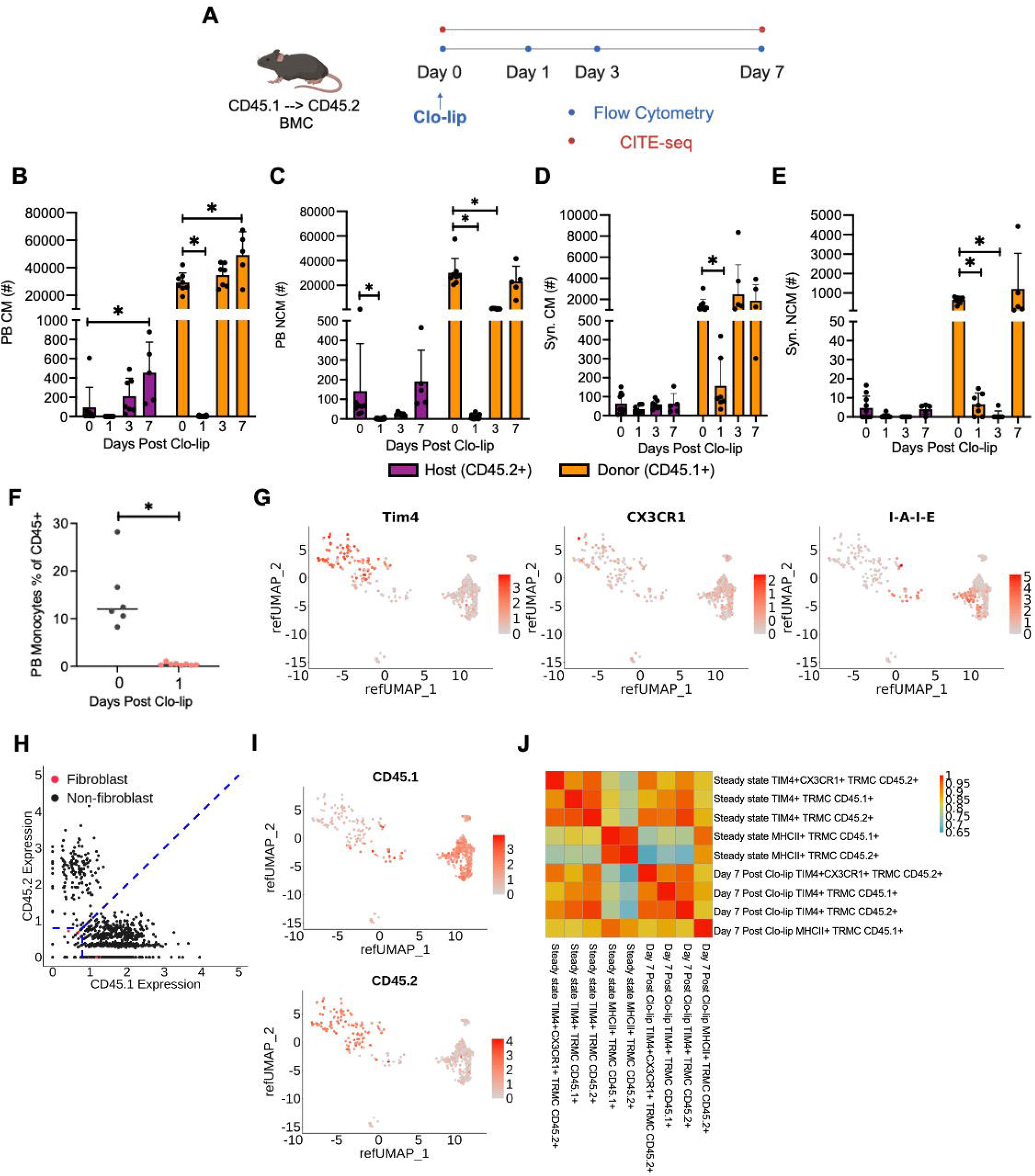
Clo-lip depleted monocytes in PB and syn. tissue as expected. (A) Schematic experimental layout of performing flow cytometry and CITE-seq on CD45.1→CD45.2 B6 BMC mice following the administration of Clo-lip. (B) Numbers of PB host-derived and donor derived CM on day 0, 1, 3, 7 post Clo-lip administration (n ≥ 5). (C) Numbers of PB host-derived and donor derived NCM on day 0, 1, 3, 7 post Clo-lip administration (n ≥ 5). (D) Numbers of Syn. host-derived and donor derived CM on day 0, 1, 3, 7 post Clo-lip administration (n ≥ 5). (E) Numbers of Syn. host-derived and donor derived NCM on day 0, 1, 3, 7 post Clo-lip administration (n ≥ 5). (F) Percents of monocyte of CD45^+^ cells in steady state BMC and day 1 post Clo-lip administration (n ≥ 6). (G) Feature plots of the expression of Tim4, CX3CR1, and I-A-I-E (ADT surface markers). (H) Feature scatter plot showing the classification of CD45.1^+^ and CD45.2^+^ cells. (I) Feature plots of the expression of CD45.1 and CD45.2 (ADT surface marker). (J) Heatmap showing the correlation of the three TRMC subsets split by CD45 expression from the steady state and day 7 post Clo-lip BMC datasets. All graphs were displayed as mean ± SD. P value was calculated by unpaired t-test with * p < 0.05

**Supplemental Figure 4.**
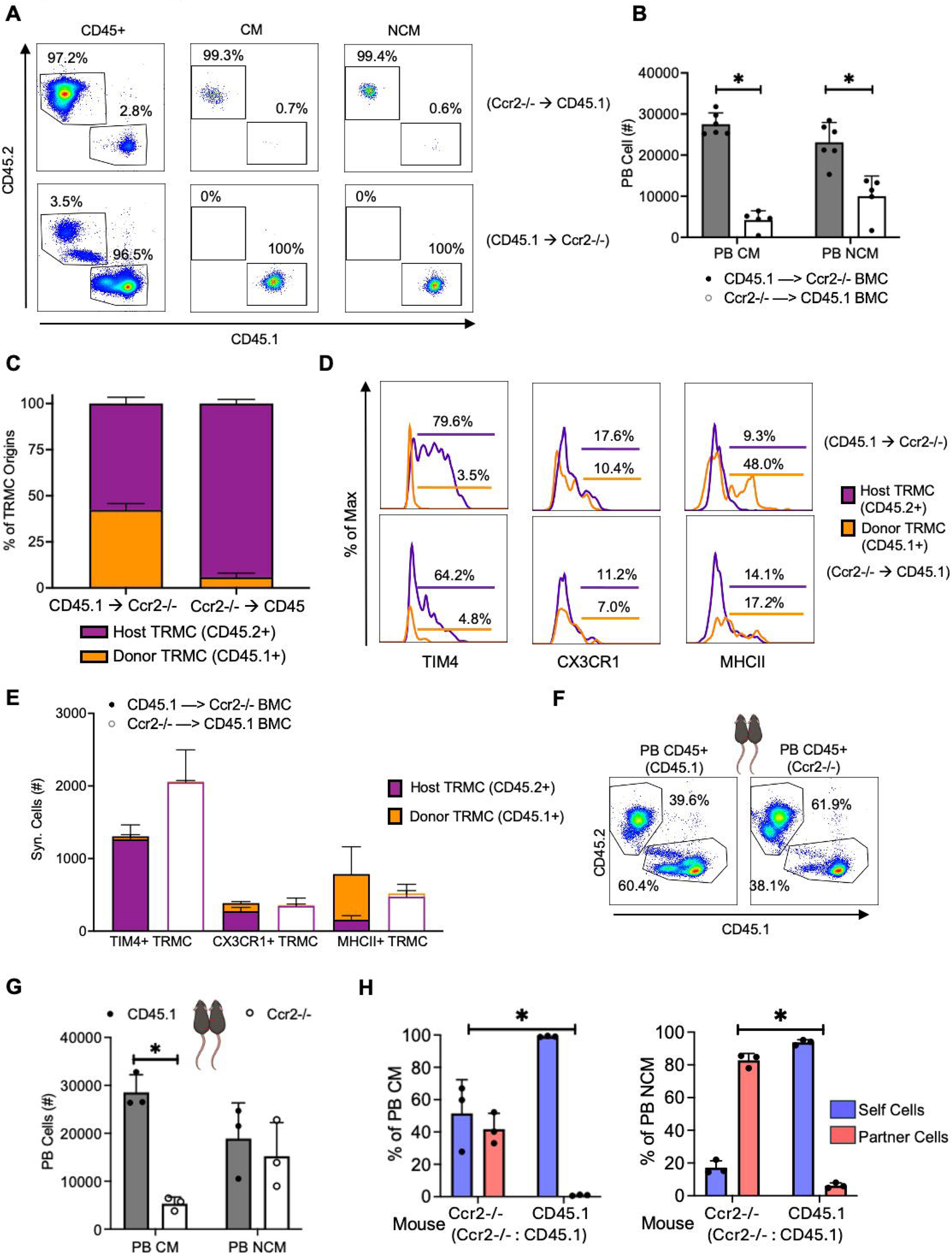
Host-derived TRMC compensate for the reduced donor-derived MHCII^+^ TRMC due to Ccr2 deficiency. (A) Percents of CD45.1+ and CD45.2^+^ cells in PB CD45^+^ cells, CM, and NCM in reciprocal Ccr2^−/−^ →CD45.1 (numbers are representatives of the averages of n = 4) and CD45.1→Ccr2^−/−^ BMC (numbers are representatives of the averages of n = 6). (B) Numbers of PB CM and NCM in reciprocal Ccr2^−/−^ →CD45.1 (n = 4) and CD45.1→Ccr2^−/−^ BMC (n = 6). (C) Percentages of donor-derived and host-derived TRMC in Ccr2^−/−^ →CD45.1 (n = 4) and CD45.1→Ccr2^−/−^ BMC (n = 6). (D) Expression and positive percentages of TIM4, CX3CR1, and MHCII of host-derived TRMC and donor-derived TRMC in reciprocal Ccr2^−/−^ →CD45.1 and CD45.1→Ccr2^−/−^ BMC, respectively. (E) Numbers of host-derived and donor-derived TIM4^+^, CX3CR1^+^, MHCII^+^ TRMC in reciprocal Ccr2^−/−^ →CD45.1 and CD45.1→Ccr2^−/−^ BMC, respectively. (F) Percents of CD45.1^+^ and CD45.2^+^ PB CD45^+^ cells in CD45.1:Ccr2^−/−^ parabiosis mice. (numbers are representatives of the averages of n = 3). (G) Numbers of PB CM and NCM in CD45.1:Ccr2^−/−^ parabiosis mice (n ≥ 2). (H) Percents of self and partner cells of PB CM and PB NCM in CD45.1:Ccr2^−/−^ parabiosis mice (n ≥ 2). Chi-square test was used to compare the distribution of cell origins from PB between CD45.1 B6 and Ccr2^−/−^ from CD45.1:Ccr2^−/−^ parabiosis, under the null hypothesis that the distribution of cell origins of any cell type does not different between the two mice groups in PB. All graphs were displayed as mean ± SD. P value was calculated by unpaired t-test with * p < 0.05.

**Supplemental Figure 5.**
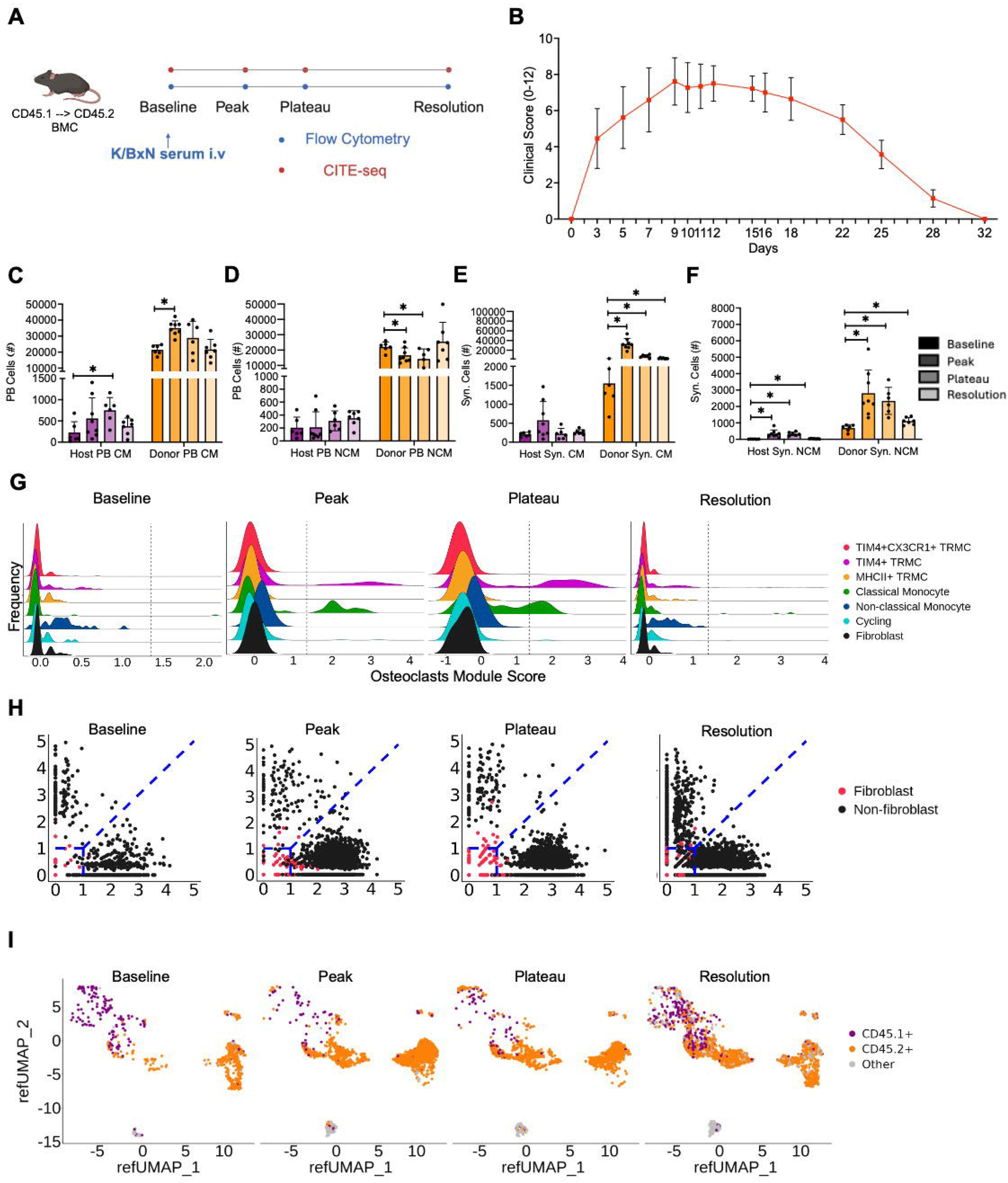
Donor-derived monocytes expanded in response to the development of inflammatory arthritis. (A) Schematic experimental layout of flow cytometry and CITE-seq on CD45.1→CD45.2 B6 BMC mice following STIA development. (B) Clinical scores of CD45.1→CD45.2 BMC mice following STIA development. (C) Numbers of PB host-derived and donor derived CM at baseline, peak, plateau, and resolution stages of STIA development (n ≥ 6). (D) Numbers of PB host-derived and donor derived NCM at baseline, peak, plateau, and resolution stages of STIA development (n ≥ 6). (E) Numbers of Syn. host-derived and donor derived CM at baseline, peak, plateau, and resolution stages of STIA development (n ≥ 5). (F) Numbers of Syn. host-derived and donor derived NCM at baseline, peak, plateau, and resolution stages of STIA development (n ≥ 5). (G) Ridge plots of osteoclast module score in seven cell populations across four STIA stages, respectively. (H) Feature scatter plot showing the classification of CD45.1^+^ and CD45.2^+^ cells across four STIA stages, respectively. (I) Annotation of CD45.1^+^ and CD45.2^+^ cells across four STIA stages, respectively. All graphs were displayed as mean ± SD. P value was calculated by unpaired t-test with * p < 0.05

**Supplemental Figure 6.**
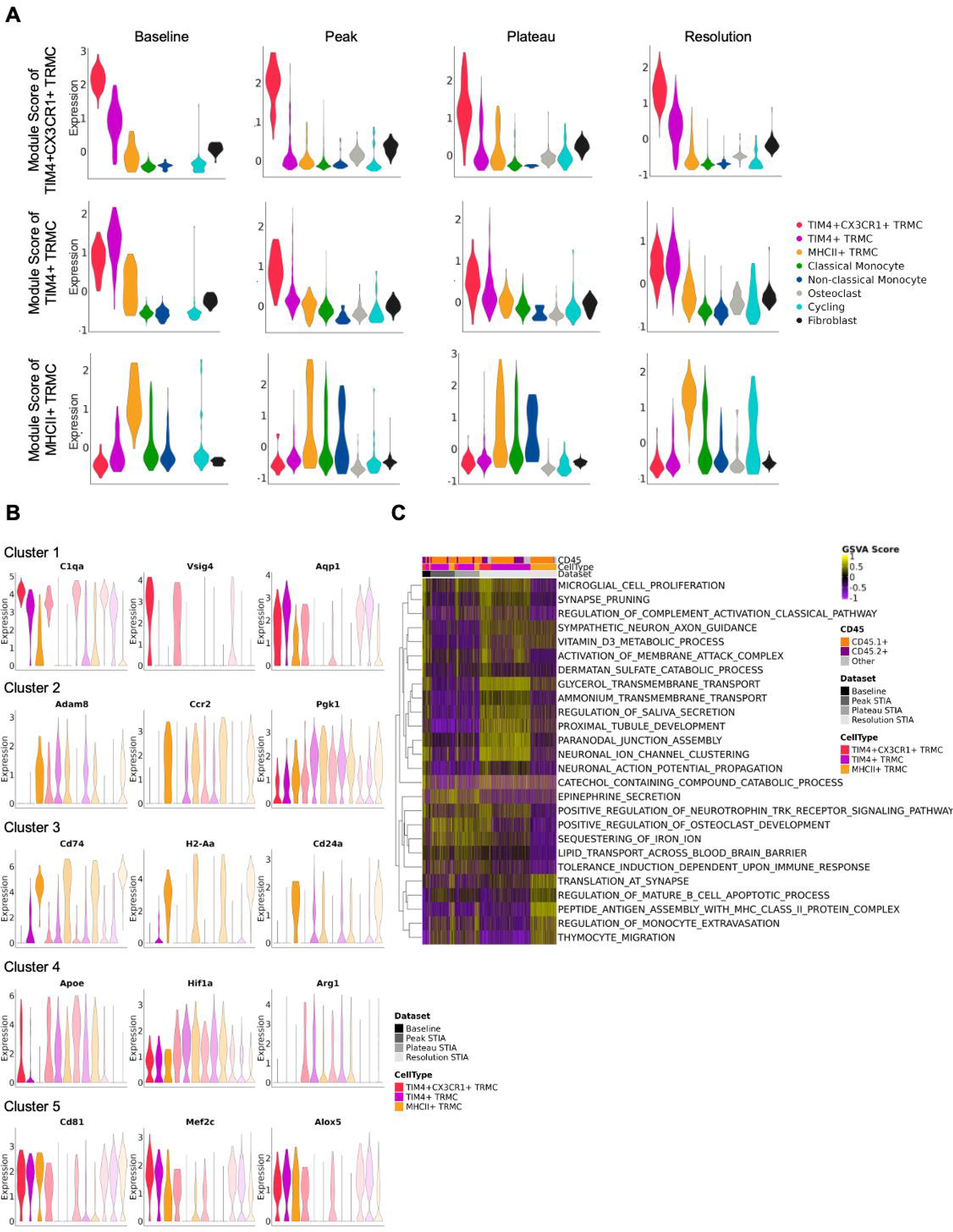
STIA introduced transcriptional changes to TRMC subsets. (A) Violin plots of the expression of the module scores of signature genes three TRMC using Fig. 2A as reference dataset across the four STIA datasets. (B) Violin plots showing the representative genes of the five clusters defined by K-means clustering of the pseudo bulk expression over 1000 variable genes of three TRMC subsets across the four STIA datasets. (C) Heatmap of gene set variation analysis (GSVA) results of the three TRMC subsets across the four stages of STIA development.

## Reference

1. McGarry, T., Hanlon, M.M., Marzaioli, V., Cunningham, C.C., Krishna, V., Murray, K., Hurson, C., Gallagher, P., Nagpal, S., Veale, D.J., and Fearon, U. (2021). Rheumatoid arthritis CD14+ monocytes display metabolic and inflammatory dysfunction, a phenotype that precedes clinical manifestation of disease. Clinical & Translational Immunology 10, e1237. 10.1002/cti2.1237.

2. Schett, G., and Gravallese, E. (2012). Bone erosion in rheumatoid arthritis: mechanisms, diagnosis and treatment. Nat Rev Rheumatol 8, 656–664. 10.1038/nrrheum.2012.153.

3. Pap, T., and Korb-Pap, A. (2015). Cartilage damage in osteoarthritis and rheumatoid arthritis—two unequal siblings. Nature Reviews Rheumatology 11, 606–615. 10.1038/nrrheum.2015.95.

4. Guo, Q., Wang, Y., Xu, D., Nossent, J., Pavlos, N.J., and Xu, J. (2018). Rheumatoid arthritis: pathological mechanisms and modern pharmacologic therapies. Bone Research 6, 15. 10.1038/s41413-018-0016-9.

5. Korganow, A.-S., Ji, H., Mangialaio, S., Duchatelle, V., Pelanda, R., Martin, T., Degott, C., Kikutani, H., Rajewsky, K., Pasquali, J.-L., et al. (1999). From Systemic T Cell Self-Reactivity to Organ-Specific Autoimmune Disease via Immunoglobulins. Immunity 10, 451–461. 10.1016/S1074-7613(00)80045-X.

6. Montgomery, A.B., Chen, S.Y., Wang, Y., Gadhvi, G., Mayr, M.G., Cuda, C.M., Dominguez, S., Moradeke Makinde, H.K., Gurra, M.G., Misharin, A.V., et al. (2023). Tissue-resident, extravascular Ly6c(-) monocytes are critical for inflammation in the synovium. Cell Rep 42, 112513. 10.1016/j.celrep.2023.112513.

7. Misharin, A.V., Cuda, C.M., Saber, R., Turner, J.D., Gierut, A.K., Haines, G.K., 3rd, Berdnikovs, S., Filer, A., Clark, A.R., Buckley, C.D., et al. (2014). Nonclassical Ly6C(-) monocytes drive the development of inflammatory arthritis in mice. Cell Rep 9, 591–604. 10.1016/j.celrep.2014.09.032.

8. Culemann, S., Grüneboom, A., Nicolás-Ávila, J.Á., Weidner, D., Lämmle, K.F., Rothe, T., Quintana, J.A., Kirchner, P., Krljanac, B., Eberhardt, M., et al. (2019). Locally renewing resident synovial macrophages provide a protective barrier for the joint. Nature 572, 670–675. 10.1038/s41586-019-1471-1.

9. Geissmann, F., Jung, S., and Littman, D.R. (2003). Blood monocytes consist of two principal subsets with distinct migratory properties. Immunity 19, 71–82. 10.1016/s1074-7613(03)00174-2.

10. Guilliams, M., Mildner, A., and Yona, S. (2018). Developmental and Functional Heterogeneity of Monocytes. 49, 595–613.

11. Jung, S., Aliberti, J., Graemmel, P., Sunshine, M.J., Kreutzberg, G.W., Sher, A., and Littman, D.R. (2000). Analysis of fractalkine receptor CX(3)CR1 function by targeted deletion and green fluorescent protein reporter gene insertion. Molecular and Cellular Biology 20, 4106–4114.

12. Silva, H.M., Báfica, A., Rodrigues-Luiz, G.F., Chi, J., Santos, P.d.E.A., Reis, B.S., Hoytema van Konijnenburg, D.P., Crane, A., Arifa, R.D.N., Martin, P., et al. (2019). Vasculature-associated fat macrophages readily adapt to inflammatory and metabolic challenges. Journal of Experimental Medicine 216, 786–806. 10.1084/jem.20181049.

13. Schyns, J., Bai, Q., Ruscitti, C., Radermecker, C., De Schepper, S., Chakarov, S., Farnir, F., Pirottin, D., Ginhoux, F., Boeckxstaens, G., et al. (2019). Non-classical tissue monocytes and two functionally distinct populations of interstitial macrophages populate the mouse lung. Nature Communications 10, 3964. 10.1038/s41467-019-11843-0.

14. Liu, X., Ren, Z., Tan, C., Núñez-Santana, F.L., Kelly, M.E., Yan, Y., Sun, H., Abdala-Valencia, H., Yang, W., Wu, Q., et al. (2024). Inducible CCR2+ nonclassical monocytes mediate the regression of cancer metastasis. J Clin Invest 134. 10.1172/jci179527.

15. Kamran, P., Sereti, K.I., Zhao, P., Ali, S.R., Weissman, I.L., and Ardehali, R. (2013). Parabiosis in mice: a detailed protocol. J Vis Exp. 10.3791/50556.

16. Wang, Y., Dowling, S.D., Rodriguez, V., Maciuch, J., Mayer, M., Therron, T., Shaw, T.N., Gurra, M.G., Shah, C.L., Makinde, H.-K.M., et al. (2025). Comprehensive analysis of myeloid reporter mice. bioRxiv, 2025.2002.2024.639159. 10.1101/2025.02.24.639159.

17. Shaw, T.N., Houston, S.A., Wemyss, K., Bridgeman, H.M., Barbera, T.A., Zangerle-Murray, T., Strangward, P., Ridley, A.J.L., Wang, P., Tamoutounour, S., et al. (2018). Tissue-resident macrophages in the intestine are long lived and defined by Tim-4 and CD4 expression. Journal of Experimental Medicine 215, 1507–1518. 10.1084/jem.20180019.

18. Etzerodt, A., Moulin, M., Doktor, T.K., Delfini, M., Mossadegh-Keller, N., Bajenoff, M., Sieweke, M.H., Moestrup, S.K., Auphan-Anezin, N., and Lawrence, T. (2020). Tissue-resident macrophages in omentum promote metastatic spread of ovarian cancer. Journal of Experimental Medicine 217. 10.1084/jem.20191869.

19. Delany, A.M., and Hankenson, K.D. (2009). Thrombospondin-2 and SPARC/osteonectin are critical regulators of bone remodeling. J Cell Commun Signal 3, 227–238 10.1007/s12079-009-0076-0.

20. Schulte, S., Podlog, L.W., Hamson-Utley, J.J., Strathmann, F.G., and Strüder, H.K. (2014). A systematic review of the biomarker S100B: implications for sport-related concussion management. J Athl Train 49, 830–850 10.4085/1062-6050-49.3.33.

21. Pixley, F.J. (2012). Macrophage Migration and Its Regulation by CSF-1. Int J Cell Biol 2012, 501962 10.1155/2012/501962.

22. Gschwandtner, M., Derler, R., and Midwood, K.S. (2019). More Than Just Attractive: How CCL2 Influences Myeloid Cell Behavior Beyond Chemotaxis. Front Immunol 10, 2759. 10.3389/fimmu.2019.02759.

23. Zlotnik, A., and Yoshie, O. (2012). The chemokine superfamily revisited. Immunity 36, 705–716. 10.1016/j.immuni.2012.05.008.

24. Zhang, Y., Cai, H., Liao, Y., Zhu, Y., Wang, F., and Hou, J. (2020). Activation of PGK1 under hypoxic conditions promotes glycolysis and increases stem cell-like properties and the epithelial-mesenchymal transition in oral squamous cell carcinoma cells via the AKT signalling pathway. Int J Oncol 57, 743–755. 10.3892/ijo.2020.5083.

25. Bernstein, B.E., and Hol, W.G. (1998). Crystal structures of substrates and products bound to the phosphoglycerate kinase active site reveal the catalytic mechanism. Biochemistry 37, 4429–4436. 10.1021/bi9724117.

26. Kuziel, W.A., Morgan, S.J., Dawson, T.C., Griffin, S., Smithies, O., Ley, K., and Maeda, N. (1997). Severe reduction in leukocyte adhesion and monocyte extravasation in mice deficient in CC chemokine receptor 2. Proc Natl Acad Sci U S A 94, 12053–12058. 10.1073/pnas.94.22.12053.

27. Hajal, C., Shin, Y., Li, L., Serrano, J.C., Jacks, T., and Kamm, R.D. The CCL2-CCR2 astrocyte-cancer cell axis in tumor extravasation at the brain. Science Advances 7, eabg8139. 10.1126/sciadv.abg8139.

28. Yona, S., Kim, K.W., Wolf, Y., Mildner, A., Varol, D., Breker, M., Strauss-Ayali, D., Viukov, S., Guilliams, M., Misharin, A., et al. (2013). Fate mapping reveals origins and dynamics of monocytes and tissue macrophages under homeostasis. Immunity 38, 79–91. 10.1016/j.immuni.2012.12.001.

29. Ginhoux, F., Greter, M., Leboeuf, M., Nandi, S., See, P., Gokhan, S., Mehler, M.F., Conway, S.J., Ng, L.G., Stanley, E.R., et al. (2010). Fate Mapping Analysis Reveals That Adult Microglia Derive from Primitive Macrophages. Science 330, 841–845. 10.1126/science.1194637.

30. Miyamoto, Y., Kikuta, J., Matsui, T., Hasegawa, T., Fujii, K., Okuzaki, D., Liu, Y.-c., Yoshioka, T., Seno, S., Motooka, D., et al. (2024). Periportal macrophages protect against commensal-driven liver inflammation. Nature 629, 901–909. 10.1038/s41586-024-07372-6.

31. Wang, P.L., Yim, A.K.Y., Kim, K.-W., Avey, D., Czepielewski, R.S., Colonna, M., Milbrandt, J., and Randolph, G.J. (2020). Peripheral nerve resident macrophages share tissue-specific programming and features of activated microglia. Nature Communications 11, 2552. 10.1038/s41467-020-16355-w.

32. Boesch, M., Lindhorst, A., Feio-Azevedo, R., Brescia, P., Silvestri, A., Lannoo, M., Deleus, E., Jaekers, J., Topal, H., Topal, B., et al. (2024). Adipose tissue macrophage dysfunction is associated with a breach of vascular integrity in NASH. Journal of Hepatology 80, 397–408. 10.1016/j.jhep.2023.10.039.

33. Alivernini, S., MacDonald, L., Elmesmari, A., Finlay, S., Tolusso, B., Gigante, M.R., Petricca, L., Di Mario, C., Bui, L., Perniola, S., et al. (2020). Distinct synovial tissue macrophage subsets regulate inflammation and remission in rheumatoid arthritis. Nature Medicine 26, 1295–1306. 10.1038/s41591-020-0939-8.

34. Zhang, F., Wei, K., Slowikowski, K., Fonseka, C.Y., Rao, D.A., Kelly, S., Goodman, S.M., Tabechian, D., Hughes, L.B., Salomon-Escoto, K., et al. (2019). Defining inflammatory cell states in rheumatoid arthritis joint synovial tissues by integrating single-cell transcriptomics and mass cytometry. Nature Immunology 20, 928–942. 10.1038/s41590-019-0378-1.

35. Schulz, C., Gomez Perdiguero, E., Chorro, L., Szabo-Rogers, H., Cagnard, N., Kierdorf, K., Prinz, M., Wu, B., Jacobsen, S.E., Pollard, J.W., et al. (2012). A lineage of myeloid cells independent of Myb and hematopoietic stem cells. Science 336, 86–90. 10.1126/science.1219179.

36. Rajan, W.D., Wojtas, B., Gielniewski, B., Miró-Mur, F., Pedragosa, J., Zawadzka, M., Pilanc, P., Planas, A.M., and Kaminska, B. (2020). Defining molecular identity and fates of CNS-border associated macrophages after ischemic stroke in rodents and humans. Neurobiology of Disease 137, 104722. 10.1016/j.nbd.2019.104722.

37. Li, Q., and Barres, B.A. (2018). Microglia and macrophages in brain homeostasis and disease. Nature Reviews Immunology 18, 225–242. 10.1038/nri.2017.125.

38. Goldmann, T., Wieghofer, P., Jordão, M.J.C., Prutek, F., Hagemeyer, N., Frenzel, K., Amann, L., Staszewski, O., Kierdorf, K., Krueger, M., et al. (2016). Origin, fate and dynamics of macrophages at central nervous system interfaces. Nature Immunology 17, 797–805. 10.1038/ni.3423.

39. Dermitzakis, I., Theotokis, P., Evangelidis, P., Delilampou, E., Evangelidis, N., Chatzisavvidou, A., Avramidou, E., and Manthou, M.E. (2023). CNS Border-Associated Macrophages: Ontogeny and Potential Implication in Disease. Curr Issues Mol Biol 45, 4285–4300 10.3390/cimb45050272.

40. Gerganova, G., Riddell, A., and Miller, A.A. (2022). CNS border-associated macrophages in the homeostatic and ischaemic brain. Pharmacology & Therapeutics 240, 108220. 10.1016/j.pharmthera.2022.108220.

41. Utz, S.G., See, P., Mildenberger, W., Thion, M.S., Silvin, A., Lutz, M., Ingelfinger, F., Rayan, N.A., Lelios, I., Buttgereit, A., et al. (2020). Early Fate Defines Microglia and Non-parenchymal Brain Macrophage Development. Cell 181, 557–573.e518. 10.1016/j.cell.2020.03.021.

42. Boring, L., Gosling, J., Chensue, S.W., Kunkel, S.L., Farese, R.V., Jr., Broxmeyer, H.E., and Charo, I.F. (1997). Impaired monocyte migration and reduced type 1 (Th1) cytokine responses in C-C chemokine receptor 2 knockout mice. J Clin Invest 100, 2552–2561 10.1172/jci119798.

43. Tsou, C.L., Peters, W., Si, Y., Slaymaker, S., Aslanian, A.M., Weisberg, S.P., Mack, M., and Charo, I.F. (2007). Critical roles for CCR2 and MCP-3 in monocyte mobilization from bone marrow and recruitment to inflammatory sites. J Clin Invest 117, 902–909. 10.1172/jci29919.

44. Puchner, A., Saferding, V., Bonelli, M., Mikami, Y., Hofmann, M., Brunner, J.S., Caldera, M., Goncalves-Alves, E., Binder, N.B., Fischer, A., et al. (2018). Non-classical monocytes as mediators of tissue destruction in arthritis. Ann Rheum Dis 77, 1490–1497. 10.1136/annrheumdis-2018-213250.

45. Brühl, H., Cihak, J., Plachý, J., Kunz-Schughart, L., Niedermeier, M., Denzel, A., Rodriguez Gomez, M., Talke, Y., Luckow, B., Stangassinger, M., and Mack, M. (2007). Targeting of Gr-1+,CCR2+ monocytes in collagen-induced arthritis. Arthritis Rheum 56, 2975–2985. 10.1002/art.22854.

46. Quinones, M.P., Ahuja, S.K., Jimenez, F., Schaefer, J., Garavito, E., Rao, A., Chenaux, G., Reddick, R.L., Kuziel, W.A., and Ahuja, S.S. (2004). Experimental arthritis in CC chemokine receptor 2-null mice closely mimics severe human rheumatoid arthritis. J Clin Invest 113, 856–866 10.1172/jci20126.

47. Kim, K.-W., Williams, J.W., Wang, Y.-T., Ivanov, S., Gilfillan, S., Colonna, M., Virgin, H.W., Gautier, E.L., and Randolph, G.J. (2016). MHC II+ resident peritoneal and pleural macrophages rely on IRF4 for development from circulating monocytes. Journal of Experimental Medicine 213, 1951–1959. 10.1084/jem.20160486.

48. Weisberg, S.P., Hunter, D., Huber, R., Lemieux, J., Slaymaker, S., Vaddi, K., Charo, I., Leibel, R.L., and Ferrante, A.W., Jr. (2006). CCR2 modulates inflammatory and metabolic effects of high-fat feeding. J Clin Invest 116, 115–124. 10.1172/jci24335.

49. Bain, C.C., Bravo-Blas, A., Scott, C.L., Gomez Perdiguero, E., Geissmann, F., Henri, S., Malissen, B., Osborne, L.C., Artis, D., and Mowat, A.M. (2014). Constant replenishment from circulating monocytes maintains the macrophage pool in the intestine of adult mice. Nature Immunology 15, 929–937. 10.1038/ni.2967.

50. Raghu, H., Lepus, C.M., Wang, Q., Wong, H.H., Lingampalli, N., Oliviero, F., Punzi, L., Giori, N.J., Goodman, S.B., Chu, C.R., et al. (2017). CCL2/CCR2, but not CCL5/CCR5, mediates monocyte recruitment, inflammation and cartilage destruction in osteoarthritis. Ann Rheum Dis 76, 914–922. 10.1136/annrheumdis-2016-210426.

51. Kadl, A., Galkina, E., and Leitinger, N. (2009). Induction of CCR2-dependent macrophage accumulation by oxidized phospholipids in the air-pouch model of inflammation. Arthritis Rheum 60, 1362–1371. 10.1002/art.24448.

52. Xie, Q., Klesney-Tait, J., Keck, K., Parlet, C., Borcherding, N., Kolb, R., Li, W., Tygrett, L., Waldschmidt, T., Olivier, A., et al. (2014). Characterization of a novel mouse model with genetic deletion of CD177. Protein & Cell 6, 117–126. 10.1007/s13238-014-0109-1.

